# Human basal radial glia morphotypes are transcriptionally distinct and exhibit different cell fate determination

**DOI:** 10.64898/2026.01.22.701090

**Authors:** Flaminia Kaluthantrige Don, Carlotta Barelli, Matteo Bonfanti, Michela Passi, Roberta Bosotti, Luca Wagner, Emanuele Capra, Ilaria Bertani, Dario Ricca, Francesca Casagrande, Alessandra Fasciani, Clelia Peano, Nereo Kalebic

## Abstract

Basal radial glia (bRG) are key neural progenitors driving human neocortical expansion. They exhibit remarkable morphological heterogeneity, yet the stability and functional significance of their distinct morphotypes remains unclear. Using human cortical brain organoids combined with long-term live imaging and morphology-resolved spatial transcriptomics (CellShape-seq), we show that bRG morphotypes display distinct morphodynamic behaviors, proliferative capacities and transcriptional profiles. While bifurcated bRG remodel extensively during mitosis to produce morphologically diverse progeny, multipolar cells are most morphologically flexible during interphase. Multipolar bRG further show the greatest proliferative capacity and the transcriptional signature related to progenitor state. Bifurcated bRG are least proliferative and are enriched for the multifunctional gene expression regulator YBX1. Pharmacological inhibition of YBX1 depletes bifurcated bRG, reduces neurogenesis, and promotes glial commitment. Our findings link progenitor morphology, gene expression and fate, providing a framework for understanding the cellular logic of human cortical development.

## Introduction

The human cerebral cortex, particularly the neocortex, underlies higher cognitive functions and exhibits remarkable expansion and complexity compared to that of other mammals. This evolutionary enlargement is mostly driven by the proliferative capacity and lineage diversity of neural progenitors. Among these, basal progenitors (BPs), play a central role ^1–9^. BPs encompass basal intermediate progenitors (bIPs), which are more restricted and typically neurogenic, and basal (or outer) radial glia (bRG/oRG), which possess enhanced self-renewal capacity ^10^. Comparative analyses across rodents, carnivores, non-human primates and humans have shown that neocortical expansion correlates with increased abundance, proliferative capacity and molecular diversification of BPs, in particular bRG ^1–3,7,11,12^.

bRG are characterized by their distinct morphological features, including long basal processes that often span multiple histological zones of the developing cortical wall ^13–15^. These cellular protrusions mediate interactions with diverse extracellular environments and provide structural scaffold for migrating neurons, positioning bRG as key organizers of cortical architecture ^7,16^. bRG self-renewal capacity and their structural specialization make these progenitors key players in supporting the massive increase in neuron number and cortical surface area in humans.

Recent studies have revealed that bRG are morphologically heterogeneous, with morphological subtypes (morphotypes) differing in process number, orientation, and spatial positioning within the germinal zones ^17–21^. Such diversity is thought to reflect functional specializations, with cellular protrusions allowing access to niche-derived factors, mechanical cues, and polarity signals ^7,22^. These extrinsic factors in turn shape bRG behaviors, their proliferative capacity, and influence neuronal and glial differentiation during brain development ^7,16,23^. However, it remains unclear whether the diverse bRG morphotypes represent intrinsically distinct progenitor identities or instead reflect transient states adopted along shared developmental trajectories. Clarifying the stability and reversibility of these morphotypes is critical for understanding how the progenitor pool organizes itself during cortical expansion. Equally unresolved are the molecular programs that underlie morphological diversity as well as the link between bRG morphology and lineage competence.

To address these questions, we leveraged human cortical brain organoids (CBOs) ^24–32^, which recapitulate key features of human corticogenesis, including bRG emergence and morphological diversity. We combined long-term live imaging with morphology-resolved spatial transcriptomics (CellShape-Seq) ^33^ to link morphological dynamics and gene expression across bRG morphotypes. This multimodal approach revealed distinct morphodynamic and transcriptional signatures among morphotypes and identified YBX1 as a molecular regulator enriched in bifurcated bRG, coupling morphology with fate determination. Together, these findings establish a framework for understanding how morphological diversity encodes lineage potential and contributes to the cellular logic of human cortical development.

## Results

### Human cortical brain organoids contain the morphological heterogeneity of bRG

We employed sliced human cortical brain organoids (CBOs) ^34^ derived from the embryonic pluripotent stem cell line H9. This system was previously shown to generate abundant bRG that recapitulate the transcriptional heterogeneity of the fetal human brain ^21,34,35^. To identify *bona fide* bRG by microscopy, we focused on HOPX+ SOX2+ cells within the subventricular zone (SVZ) of CBOs. This strategy is supported by multiple *in vivo* and CBO studies demonstrating that subventricular HOPX is a robust and conserved marker of self-renewing bRG in the human neocortex ^18,36,37^.

Immunofluorescence analysis across developmental stages identified the emergence of HOPX+ SOX2+ bRG in the SVZ at week 11 CBOs, which coincides with the SVZ expansion (Figure 1A). Additionally, these bRG also expressed radial glia marker nestin (Figure S1A, B). At the same developmental stage (week 11) we further detected the emergence of late-born SATB2+ neurons (Figure S1C, D). This is consistent with the known contribution of bRG (and late bIPs) to the production of late-born neurons ^38–40^.

**Figure 1.**
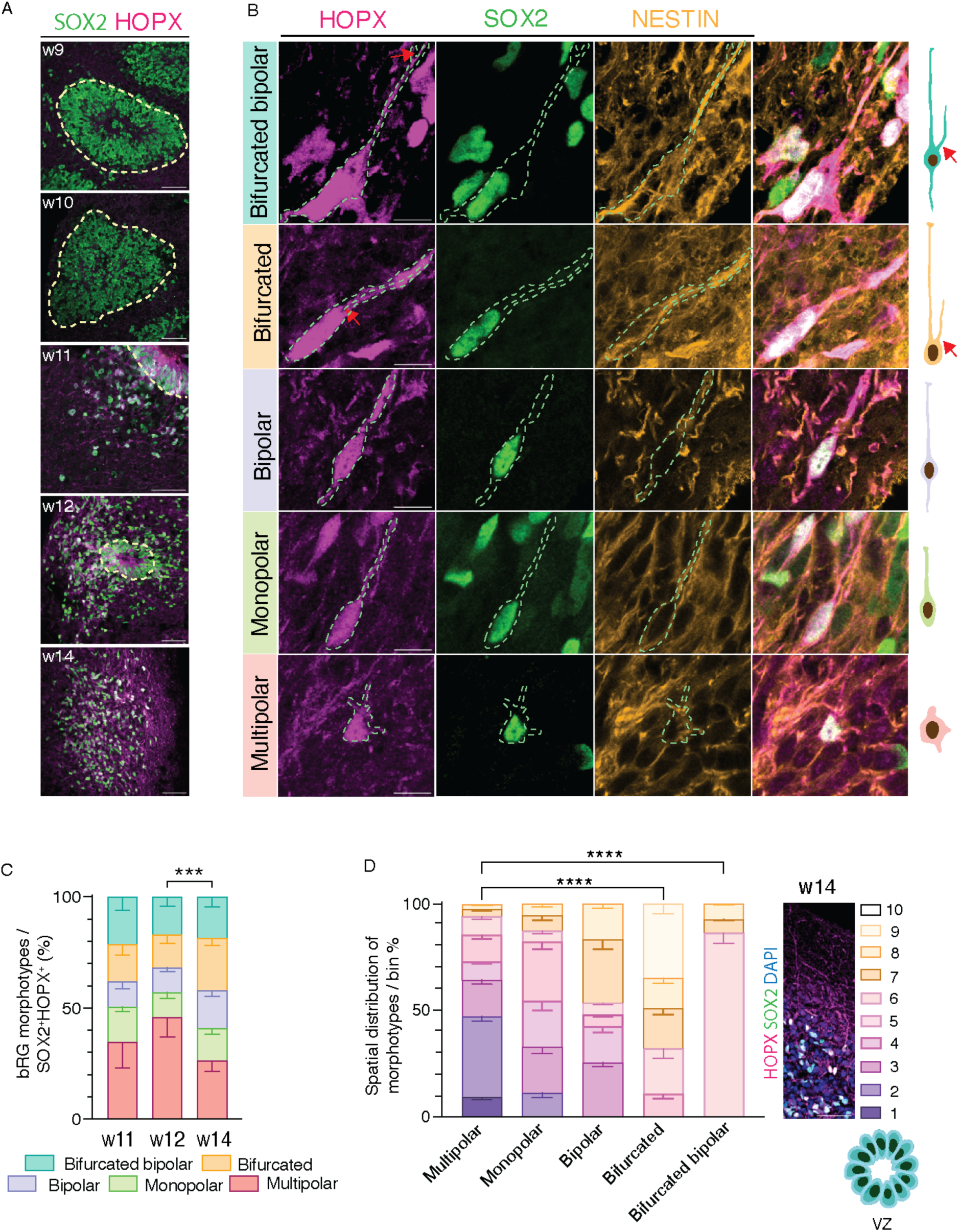
Cortical brain organoids (CBOs) contain the morphological heterogeneity of bRG. **A)** Immunofluorescence (IF) for SOX2 and HOPX of week (w)9 to w14 CBOs. Dashed yellow line marks the basal side of the ventricular zone (VZ). Scale bar, 50 µm**. B)** IF for SOX2, HOPX and NESTIN of the five individual bRG morphotypes (listed on the left) in CBOs along with the schematic representation of morphotypes (right). Dashed line, cell outline to mark the morphology. Red arrow, bifurcation. Scale bar, 10 µm. **C)** Quantification of bRG morphotypes distribution in CBOs at w11, 12, 14. Data represent 10 (w11, 12) or 14 (w14) CBOs with 471 (w11), 527 (w12) and 792 (w14) bRG. Error bars, SEM; 2-way ANOVA with Bonferroni’s post hoc test; ***p < 0.001**. D)** Quantification of the spatial distribution of bRG morphotypes along the apico-basal axis (divided in 10 bins) of w11 and w14 CBOs; N = 12 CBOs with 297 bRG. Error bars, SEM; Two-way ANOVA and Bonferroni’s post hoc test; Multipolar vs Bifurcated **** p < 0.0001; Multipolar vs Bifurcated bipolar **** p < 0.0001.

We next examined the morphology of HOPX+ SOX2+ cells using HOPX signal to trace cell shape, as previously described ^18^. In CBOs, we reproducibly identified five bRG morphotypes - multipolar, monopolar, bipolar, bifurcated, and bifurcated bipolar - distinguished by the number and orientation of processes (Figure 1B, C). This classification mirrors morphologies observed in fetal human tissue ^18^. Since CBOs contain multiple ventricle-like lumens and exhibit disrupted global apico–basal polarity, all monopolar cells with either apically or basally directed processes were grouped into a single category. While all morphotypes were represented, their relative proportions differed from those in fetal tissue, with CBOs showing an increased fraction of bifurcated bRG ^18^. Moreover, the proportion of this bRG morphotype further increased between weeks 11 and 14 (Figure 1C). Interestingly, this was paralleled with a relative increase in proportion of late-born SATB2+ neurons *versus* early-born CTIP2+ neurons (Figure S1C, D).

Mapping the spatial distribution of bRG morphotypes in CBOs revealed a distinct apico-basal gradient. Notably, multipolar cells were preferentially found in the apical-most portion of the SVZ, whereas bifurcated bipolar bRG were enriched in the basal-most part (Figure 1D). Monopolar, bipolar and bifurcated bRG were distributed sequentially along the apico-basal axis. Finally, to confirm that the multipolar cells were not basal intermediate progenitors (bIPs), we stained for TBR2 ^41^, whose absence excluded bIPs (Figure S2).

Taken together, these data show that bRG morphotypes emerge contemporarily in CBOs, but they are differentially distributed along the apico-basal axis. Furthermore, our data indicate that human CBOs contain a morphological heterogeneity of bRG similar to the one observed in the developing human neocortex ^18^ and can thus be regarded as a valid system to dissect bRG morphodynamic behavior and gene expression.

### bRG-enriched BP population shows morphological inheritance and flexibility

We next used CBOs to examine the stability and dynamics of bRG morphotypes over time. We previously hypothesized that bRG might inherit the morphotype status from the mother cell yet also exhibit morphological flexibility to generate the full diversity observed *in vivo* ^22^. To test this, we examined bRG morphodynamics using long-term time-lapse microscopy.

Sliced 12–14-week-old CBOs were transfected with GFP and cultured for two days followed by live imaging for up to four days using confocal microscopy (Figure 2A). We quantified that >70% of GFP+ cells in the SVZ were bRG (Figure S3). While bRG normally account for around 50% BPs in human ^18^, the method of electroporation along the CBO lumens likely leads to an enrichment in bRG, as it labels aRG and their progeny which is preferentially bRG rather than bIPs or neurons at this developmental stage ^38^. We focused our analysis only on GFP+ cells that underwent a division during the imaging period, thereby excluding post-mitotic neurons and yielding a population of dividing BPs strongly enriched for bRG.

**Figure 2.**
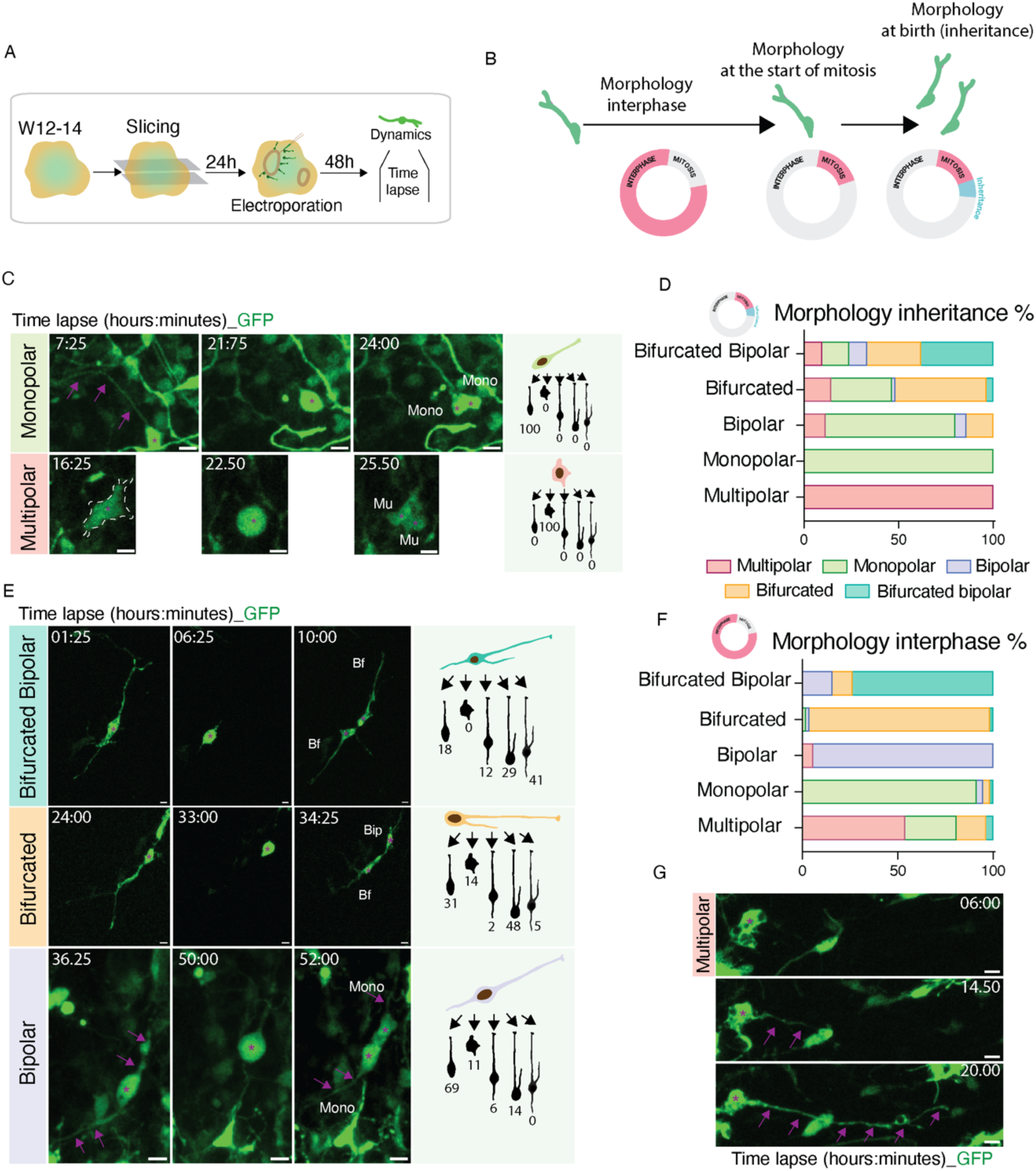
bRG-enriched BP population shows morphological inheritance and flexibility. **A)** Experimental plan of live imaging in CBOs. **B)** Schematic representation of BP morphological tracking by live imaging encompassing morphological inheritance (from mother cell to daughter cells) and morphological dynamics in interphase. **C, E)** Representative time-lapse sequences of individual BP morphotypes (listed on the left) undergoing division and the morphology of their daughter cells. Dashed line, cell morphology; Asterisk, cell body. Right, schematic representation of the morphological inheritance shown as the percentage of morphotype progeny for each mother morphotype (Bf, bifurcated; Bip, bipolar; Mono, monopolar). Number of mother cells: multipolar (10), monopolar (13), bipolar (18), bifurcated (28) and bifurcated bipolar (14). Scale bar, 10 µm. **D)** Quantification of the morphological inheritance for each morphotype. N= 144 daughter cells from 13 different time-lapse experiments. **F)** Quantification of morphological changes in interphase for each morphotype; color coding as in (D). N= 139 cells from 13 different time-lapse experiments. **G)** Representative time-lapse sequence of a multipolar cell elongating in interphase. Asterisk, cell body; Arrows, growing cell process. Scale bar, 10 µm.

Because mitosis involves characteristic morphological remodeling, we first compared the morphology of mother and daughter cells (Figure 2B). Strikingly, monopolar and multipolar cells showed complete inheritance of morphology, with 100% of daughter cells retaining the mother’s morphotype, indicating a high degree of intrinsic stability (Figure 2C, D, S4A). In contrast, bipolar, bifurcated, and bifurcated bipolar cells exhibited considerable morphological plasticity during mitosis. In particular, bifurcated and bifurcated bipolar bRG give rise to daughters spanning the full spectrum of morphotypes, thereby contributing disproportionately to the overall morphological diversity (Figure 2D, E, S4A). Similarly, bipolar cells generated a wide range of daughter cell morphologies, giving rise to all morphotypes but one (bifurcated bipolar) (Figure 2D, E, S4A).

We next assessed morphological transitions during interphase (Figure 2B). Most radial morphotypes remained remarkably stable, showing few morphological transitions (Figure 2F). In contrast, multipolar cells displayed pronounced morphological remodeling and the ability to transform into different radial morphotypes (Figure 2F, G, S4B). This suggests that multipolar cells exhibit the greatest morphological flexibility outside of mitosis.

Collectively, these findings reveal marked differences in bRG morphotypes’ propensity to morphological changes. Morphological transitions are modest during interphase, but they become markedly more frequent during mitosis (Figure S4C). Moreover, bifurcated, bipolar, and bifurcated bipolar cells contribute most to mitosis-associated morphological diversity, whereas multipolar cells remain uniquely flexible during interphase.

### bRG morphotypes exhibit different proliferative capacities

We next examined whether distinct bRG morphotypes differ in their proliferative capacity. To this end, we first performed *post hoc* cell fate analysis following live imaging (Figure 3A). After fixation and immunostaining, the progeny of imaged BPs was classified as progenitors (SOX2+) or neurons (HuC/HuD+) (Figure 3B and S5A). We were able to successfully assign identity to 48 daughter cells, representing roughly one-third of all divisions. In agreement with previous reports ^20^, most progeny were SOX2+ progenitors (Figure 3B). This suggests that a vast majority of BPs undergoes proliferative rather than differentiative divisions during the weeks 12-14 of CBO development.

**Figure 3.**
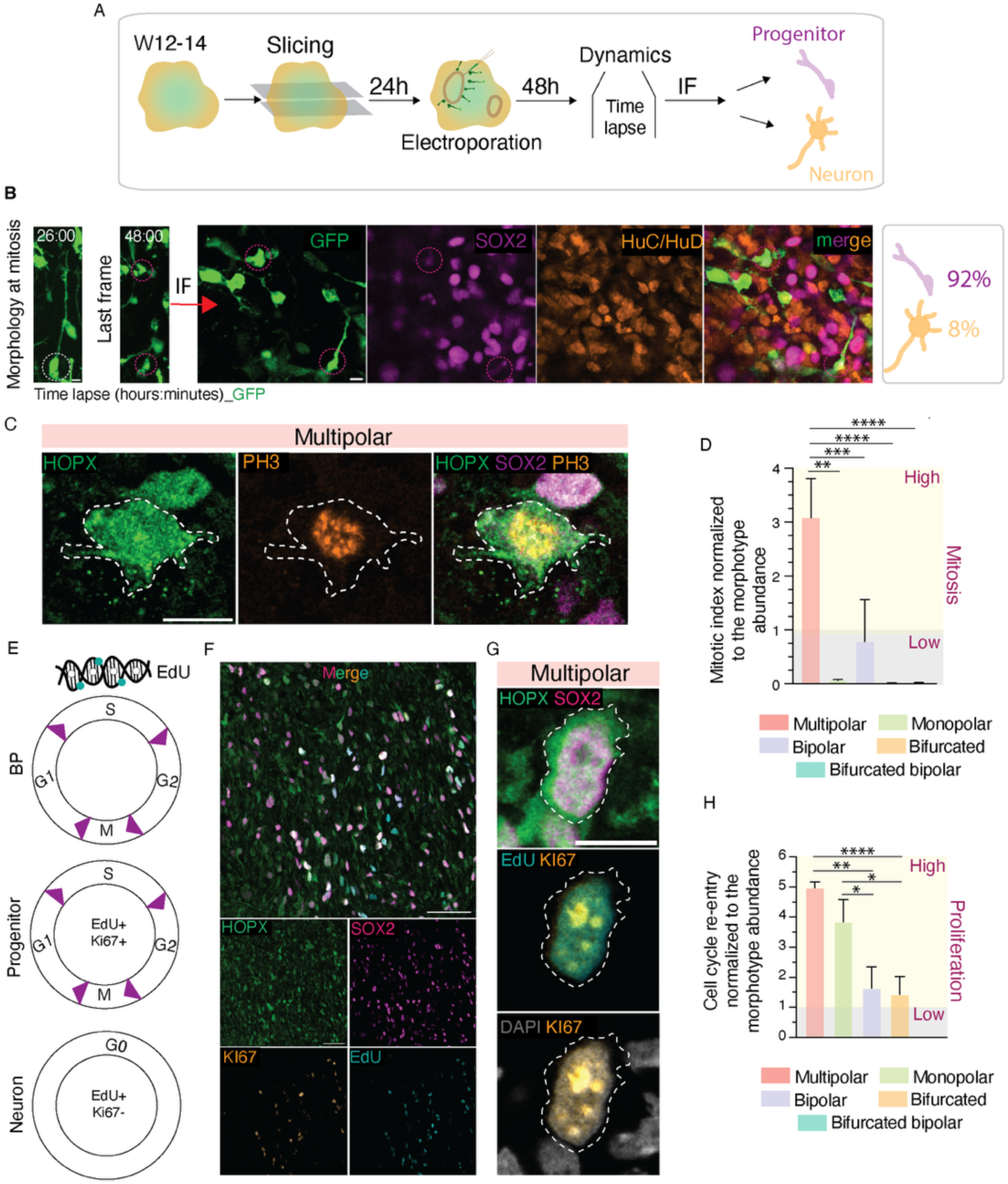
bRG morphotypes exhibit different proliferative capacities. **A)** Schematic representation of the live imaging workflow with the cell fate mapping. **B)** Quantification of bRG daughter cells (dotted line) expressing SOX2 and HuC/HuD; n = 45 fate-mapped daughter cells. Note that the vast majority of daughter cells are progenitors. Scale bar, 10 µm. **C)** Multipolar bRG expressing HOPX, SOX2 and PH3. Dashed line, cell outline. Scale bar, 10 µm. **D)** Mitotic index visualized as a fold-enrichment of PH3+ SOX2+ HOPX+ cells across morphotypes and calculated as the proportion of PH3+ SOX2+ HOPX+ cells in each morphotype normalized by the relative abundance of that morphotype. Values >1 reflect morphotypes that are disproportionately represented among the mitotic cells (High, yellow), whereas values <1 indicate under-representation (Low, grey). N= 58 bRGs; error bars, SEM; Mann-Whitney u-test, ** p< 0.01; *** p< 0.001; **** p< 0.0001. **E)** Schematic representation of the cell cycle re-entry assay. **F)** IF for EdU, KI67, SOX2 and HOPX. **G)** A cycling multipolar bRG expressing EdU+ Ki67+ SOX2+ HOPX+. **H)** Proliferative capacity visualized as a fold-enrichment of EdU+ Ki67+ SOX2+ HOPX+ cells across morphotypes and calculated as the proportion of EdU+ Ki67+ SOX2+ HOPX+ cells in each morphotype normalized by the relative abundance of that morphotype. Values >1 reflect morphotypes that are disproportionately represented among the proliferative cells (High, yellow), whereas values <1 indicate under-representation (Low, grey). N= 308 bRGs; error bars, SEM; Mann-Whitney u-test, * p< 0.05; ** p< 0.01; **** p< 0.0001.

To assess whether proliferation varies among morphotypes, we first focused on mitosis. We stained week 14 CBOs for the mitotic marker phospho-histone 3 (PH3) (Figure 3C) and quantified the mitotic index of SOX2+HOPX+ bRG (Figure S5B). Multipolar and monopolar bRG displayed a higher frequency of mitosis compared with other morphotypes. When normalized to the relative abundance of each morphotype, this difference became even more pronounced: multipolar bRG showed the greatest enrichment in mitotic activity (Figure 3D).

Such differences in mitotic frequency could be due to an increased bRG proliferative capacity or to the relative increase of time they spend in mitosis compared to the rest of the cell cycle. We hence compared cell cycle length across morphotypes and found no significant differences (Figure S5C). To measure the proliferative capacity of bRG we next performed a cell cycle re-entry assay. We treated week 14 CBOs with thymidine analogue EdU for 30 h, which corresponds to approximately 80% of the cell cycle duration (Figure S5C), allowing labelled progenitors to progress from the S-phase (when EdU is incorporated) into the next cell cycle. We then quantified the proportion of EdU+ cells that remained Ki67+ progenitors, indicating cell cycle re-entry (Figure 3E, F). Multipolar bRG exhibited the greatest cell cycle re-entry rate, followed by monopolar cells, while bipolar and bifurcated cells showed a lower proliferative capacity (Figure 3G, S5D). When normalized to the morphotype abundance, multipolar and monopolar bRG again emerged as the most proliferative (Figure 3H).

Taken together, our data indicate marked differences in proliferative capacity among bRG morphotypes. Multipolar bRG exhibit the strongest proliferative capacity, monopolar cells show intermediate and bifurcated cells are the least proliferative. Combined with the morphodynamic differences observed earlier (Figure 2), these results prompt us to examine whether specific bRG morphotypes might correspond to distinct transcriptional states as well.

### bRG morphoclasses have distinct transcriptomic signatures

To resolve the bRG transcriptome in a morphology-dependent manner, we applied our customized spatial transcriptomics pipeline, CellShape-seq, previously established on brain tumor organoids ^33^. This approach combines manual segmentation of individual cells identified by markers and morphological features with whole-transcriptome profiling using the GeoMX platform. To account for the imaging resolution of the GeoMX and increased scale of analysis, we grouped five morphotypes into three morphoclasses, similarly to what previously reported in brain tumors ^33^: (1) monopolar+bipolar morphoclass; (2) multipolar morphoclass; (3) bifurcated morphoclass (encompassing both bifurcated morphotypes) (Figure 4A). Because CellShape-seq requires FFPE material, we identified bRG within the CBOs’ SVZ as GFAP+ vimentin+ SOX2+ cells (Figure 4B), after corroborating that these cells also co-express HOPX (Figure S6). Using this strategy, we profiled 18 CBOs derived from three independent differentiations.

**Figure 4.**
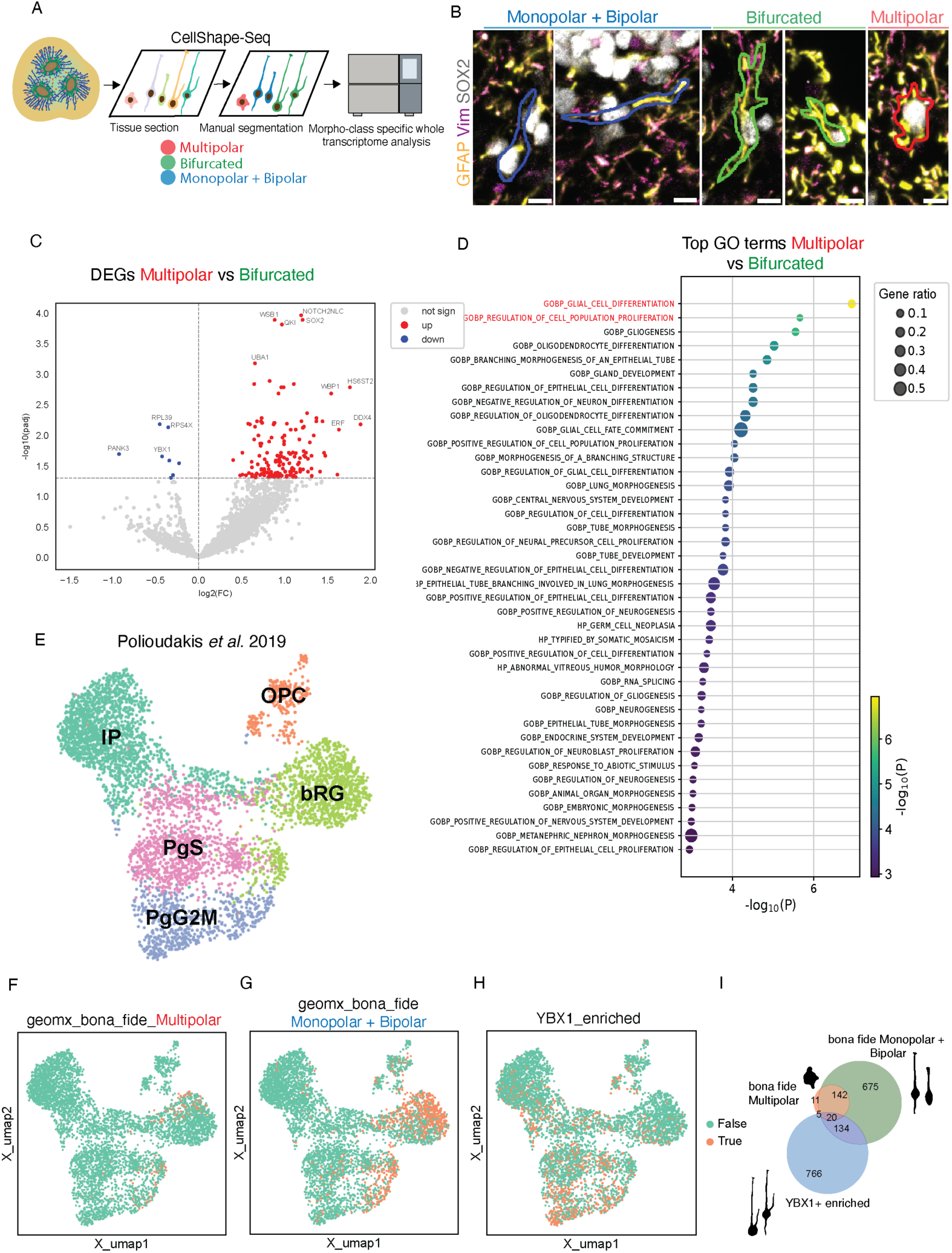
bRG morpho-classes have distinct transcriptomic signatures. **A)** Schematic representation of CellShape-seq pipeline integrating cell morphology and gene expression. For the spatial transcriptomics, bRG were grouped in 3 classes: multipolar; monopolar+bipolar; bifurcated + bifurcated bipolar. **B)** Examples of manual segmentations of the five bRG morphotypes, grouped into 3 classes and expressing GFAP, Vim and SOX2. Scale bars, 10 µm. **C)** Volcano plot showing the results of a differential expression analysis of multipolar *vs* bifurcated bRG. The x-axis, Log-Fold Change (LFC); y-axis, negative log10 of the adjusted p-value. Red dots, significantly up-regulated genes (LFC > 0 and adjusted p-value < 0.05); blue dots, significantly down-regulated genes (LFC < 0 and adjusted p-value < 0.05). **D)** List of Gene Ontology Biological Processes (GOBP) significantly upregulated in multipolar vs bifurcated bRG, selected based on a LFC < 0 and an adjusted p-value (corrected for multiple testing) < 0.05. The x-axis position and dot color indicate the significance level of pathway enrichment, while dot size represents the proportion of differentially expressed genes associated with each GO term. Top pathways pertaining to cell proliferation and glial differentiation are marked in red. **E)** Uniform Manifold Approximation and Projection (UMAP) plot of all clusters containing BPs from the single-cell RNA-seq dataset published by ^49^, defined by selecting clusters annotated as bRG (Basal Radial Glia i.e Outer Radial Glia), PgS (Progenitors in S phase), and PgG2M (Progenitors in G2/M phase), IP (Intermediate Progenitors), OPC (Oligodendrocyte Progenitor Cells). **F, G)** The UMAP representation of the progenitor cells from the ^49^ dataset showing in orange the cells identified as bona fide multipolar bRG (F) and bona fide monopolar+bipolar bRG (G). These bona fide populations were defined based on the projection of GeoMX-derived gene signatures as detailed in the Methods. **H)** The UMAP representation of the subset of progenitor cells enriched for YBX1 activity signature (see the Methods) from the ^49^ dataset. **I)** Venn diagram illustrating the overlap among the YBX1+ enriched subset (preferentially bifurcated and bifurcated bipolar bRG) with the bona fide monopolar+bipolar and multipolar bRG, as identified in (F-H) within the progenitor cells from the ^49^ dataset.

Differential gene expression analysis revealed marked transcriptomic divergence between multipolar and bifurcated bRG (Figure 4C) as well as between monopolar+bipolar and bifurcated classes (Figure S7A), but not between multipolar and monopolar+bipolar (Figure S7B). Gene Ontology (GO) analysis of multipolar *vs* bifurcated bRG highlighted enrichment for terms associated to cell proliferation and glial fate commitment in multipolar cells (Figure 4D), consistent with their higher proliferative capacity and mitotic index (Figure 3D, H). Ingenuity pathway analysis (IPA) further identified *TUBG1* (γ-tubulin) and *MOB1A* (member of the Hippo pathway) as enriched in multipolar bRG (Figure S7C) and predicted activation of upstream regulators including *TGFB1*, *EGFR*, *PDGF*, *SOX2* and *GLI1* (Figure S7D). Together, these analyses suggest a strong upregulation of neurodevelopmental pro-proliferative pathways in multipolar bRG (Figure S7E). Moreover, the analysis highlighted several genes significantly upregulated in multipolar bRG, including *SOX3*, *PAX6*, *TMX2*, *NR2F1, HGNAT* or *HS6ST2*, which are associated with intellectual disability, autism spectrum disorder and related neurodevelopmental conditions (Figure S7F).

Instead, morphoclass comprising monopolar and bipolar bRG showed fewer transcriptional differences in comparison with bifurcated bRG and none compared to multipolar bRG (Figure S7A, B). Nevertheless, the genes enriched in monopolar + bipolar *vs*. bifurcated cells were associated to cell proliferation (*TTYh1*, *FGFR3*) and neurite outgrowth (*RGMA*, *PTN*, *TTYh1*) with predicted upstream activators, RNA-binding *FMR1* and *LARP1* (Figure S8A, B), both previously implicated in neurodevelopment ^42–46^. In contrast, bifurcated morphoclass showed only a small set of upregulated genes: eight vs multipolar and five vs monopolar + bipolar (Figure 4C, S7A). Among these we identified Y-box binding protein 1 (YBX1), an RNA/DNA binding factor, and multifunctional regulator of gene expression ^47,48^.

We next mapped morphoclass-enriched transcriptomic signatures to a published human fetal brain scRNA-seq dataset ^49^. We selected five progenitor clusters, enriched for BPs and representing bRG, bIPs, oligodendrocyte progenitor cells (OPCs) and progenitors in S/G2M phases (Figure 4E). The gene expression signature of multipolar and monopolar+bipolar morphoclasses were projected onto those five clusters (Figure 4F, G). Importantly, the vast majority of the mapped cells were contained in the clusters annotated as bRG and “progenitors in S/G2M”, corroborating our microscopy-based bRG identification. Given that the signature of bifurcated bRG-enriched genes did not comprise enough genes to perform the analogous mapping, we instead utilized YBX1 as a representative marker. Based on predicted YBX1 activation through its downstream targets (Figure S8C, D), we identified YBX1-enriched progenitors across the five clusters (Figure 4H). Notably, in clusters corresponding to bRG, YBX1-enriched cells were largely non-overlapping with those mapping to multipolar or monopolar+bipolar bRG (Figure 4I). This suggests distinct transcriptional identities within the bRG lineage.

Together, these results demonstrate that bRG morphoclasses harbor distinct transcriptional programs that align with their morphological and proliferative features. Furthermore, CellShape-seq enables direct integration of morphology-resolved transcriptomics with fetal brain single-cell data, showing previously unappreciated molecular diversity within the bRG population.

### YBX1 maintains bifurcated bRG identity and restrains premature gliogenesis

Given our results that bRG morphotypes differ in morphodynamics, proliferative capacity, and gene expression signatures, we next asked whether these differences may influence downstream developmental events, such as cell fate determination. The balance between proliferation and differentiation as well as between neurogenesis and gliogenesis is a fundamental principle of cortical development and its disruption is closely linked to various neurodevelopmental disorders ^50–52^. To examine potential links between morphotype and fate, we focused on bifurcated bRG and selected *YBX1*, a gene highly enriched in this morphotype (Figure 4C). YBX1 is known to have multiple roles in controlling gene expression, from modulating mRNA stability, translation, splicing to transcriptional regulation ^47,48,53^. Additionally, during brain development, YBX1 fine-tunes spatiotemporal gene expression programs that promote progenitor self-renewal and neuronal differentiation ^54^.

We first validated YBX1 expression at the protein level in week 14 CBOs and detected YBX1 in HOPX+ SOX2+ progenitors in the SVZ (Figure S9A). To examine the functional role of YBX1, we performed pharmacological inhibition using SU056, an azopodophyllotoxin-derived small molecule, a validated and specific YBX1 inhibitor ^55^. Dose-response assays established that 100 nM and 1 μM as non-toxic and suitable concentrations for long-term treatment of CBOs (Figure S9B). Normally, YBX1 is retained in the cytoplasm, while upon stress-inducing conditions such as chemotherapy, it is translocated to the nucleus ^56,57^. In agreement with this, we detected YBX1 in the bRG cytoplasm in both the cell body and protrusions (Figure S9C), while SU056 treatment led to a nuclear translocation of YBX1 in all morphotypes (Figure S9D-F).

We used two treatment paradigms in CBOs: short-term (8 days) and long-term (30 days) exposures (Figure 5A, I). Short-term inhibition led to a pronounced relative loss of bifurcated bRGs accompanied by an apparent increase in multipolar bRG (Figure 5B, C). To examine whether such condition was due to depletion of bifurcated bRG or their transition to multipolar cells we quantified overall BP abundance and detected a strong decrease in SOX2+ progenitors in the SVZ upon YBX1 inhibition (Figure 5D and S10A). These data suggest YBX1 inhibition leads to a depletion of bifurcated bRG, while multipolar bRG, as the most proliferative morphotype (see Figure 3H), become predominant.

**Figure 5.**
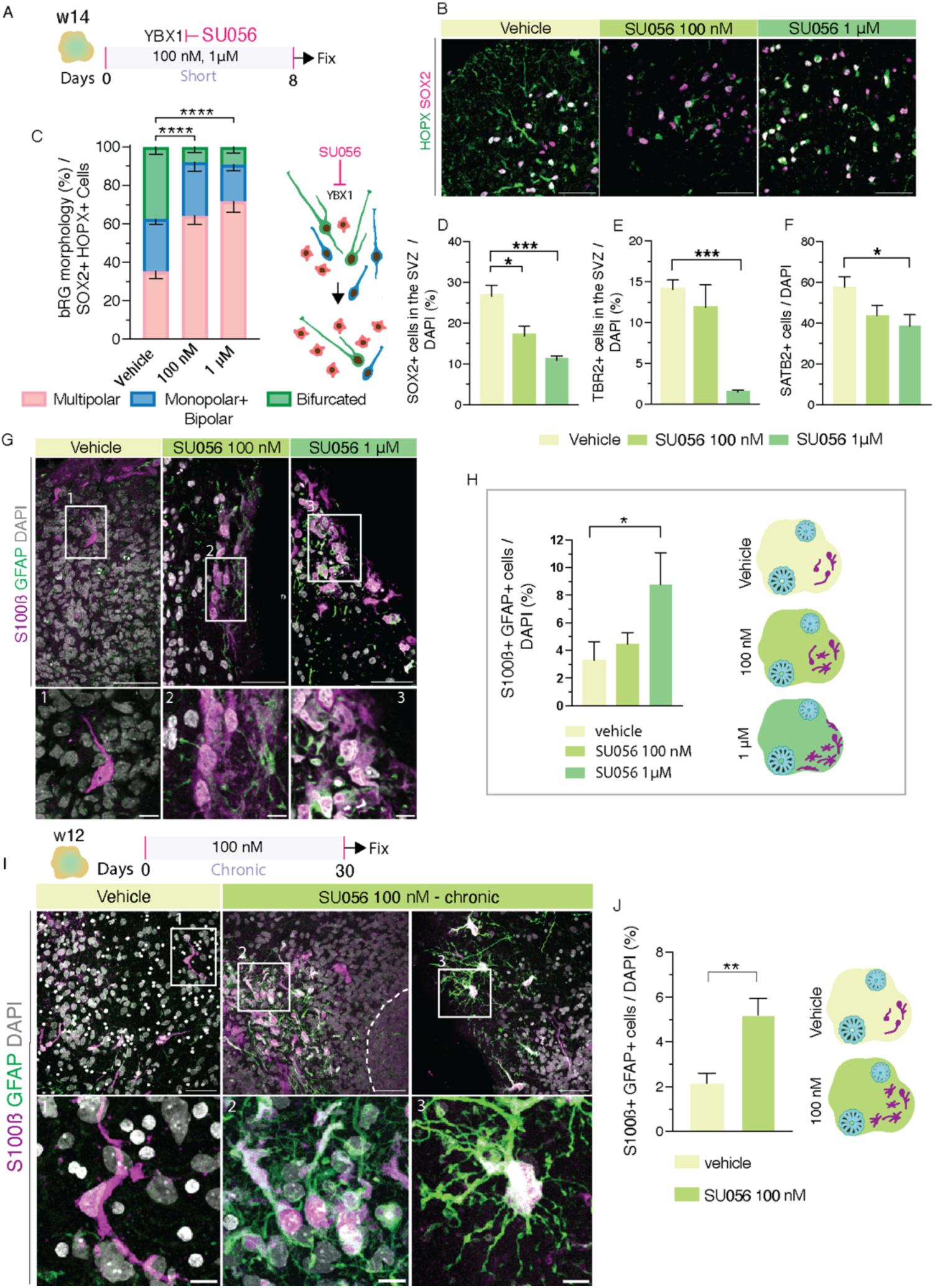
YBX1 maintains bifurcated bRG identity and restrains premature gliogenesis. **A)** Schematic representation of short-term (8 days) administration of YBX1 inhibitor SU056 in CBOs starting at w14 with two indicated concentrations. **B)** IF for HOPX and SOX2 showing bRG morphotypes upon YBX1 inhibition, with the indicated concentrations of SU056. Scale bar, 50 µm. **C)** Quantification of bRG morphotype distribution showing a reduction of bifurcated bRG upon YBX1 inhibition. N= 3 independent CBO differentiations; errors bars, SEM; Multipolar (vehicle vs SU056 100 nM, 1 μM) **** p < 0.0001; Monopolar + bipolar (vehicle vs SU056 100 nM, 1 μM) n.s.; Bifurcated (vehicle vs SU056 100 nM, 1 μM) **** p < 0.0001. **D-F)** Quantification of SOX2+ SVZ progenitors (D), TBR2+ neurogenic SVZ progenitors (E) and SATB2+ late-born neurons (F) upon short-term YBX1 inhibition in CBOs. Error bars, SEM; N= 3 independent CO differentiation; Mann-Whitney U-test, *p < 0.05, p***<0.001. **G)** IF for astrocytic markers S100ß and GFAP in CBOs in presence and absence of YBX1 inhibition with the indicated concentrations of SU056. Boxes 1-3, areas magnified below. Insets, higher magnification of astrocytes. Scale bar, 50 µm (overview); 10 µm (inset). **H)** Quantification of S100ß+ GFAP+ astrocytes, showing their increase with the high concentration of YBX1 inhibitor. Error bars, SEM; N= 3 independent CO differentiation; Mann-Whitney U-test, *p < 0.05. **I)** Long term inhibition of YBX1. Upper, schematic representation of CBOs treated for 30 days (chronic) with SU056 (w12-16). Lower, IF for astrocytic markers S100ß and GFAP in CBOs upon chronic YBX1 inhibition. Dashed line, basal limit of VZ; Boxes 1-3, areas magnified in insets below. Insets, higher magnification of astrocytes. Scale bars, 50 µm (overview); 10 µm (inset). **J)** Quantification of S100ß+ GFAP+ astrocytes, showing their increase upon chronic YBX1 inhibition. Error bars, SEM; N= 3 independent CO differentiation; Mann-Whitney u-test, ** p< 0.01.

We next investigated whether depletion of bifurcated bRG reflected premature differentiation. YBX1 inhibition markedly reduced both TBR2+ bIPs (Figure 5E and S10B) and SATB2+ late-born neurons (Figure 5F and Figure S10C). This suggests that diminished SATB2+ neuron production may result from loss of the progenitor subpopulation that generates them, i.e. bifurcated bRG. In striking contrast, YBX1 inhibition strongly enhanced glial differentiation, with a robust increase in GFAP+ S100ß+ cells in the basal compartment of the CBOs (SVZ and the CP-like structures), indicative of elevated astrocytic commitment (Figure 5G, H and Figure S10D). Indeed, these cells displayed characteristic stellate morphologies, consistent with astrocytic identity (Figure 5I and S10D). Such an increase in astrocytes was observed under both short and long treatments (Figure 5H, J) and was more prominent at 1 μM SU056 than at 100 nM (Figure 5H), suggesting a dose-dependent effect.

Together, these findings identify YBX1 as a potential regulator of bifurcated bRG state. YBX1 activity maintains the bifurcated bRG identity, and its inhibition destabilizes this morphotype, leading to a premature gliogenic commitment. Thus, YBX1 emerges as a potential molecular link coupling progenitor morphology and fate determination during human cortical development.

## Discussion

Our study links bRG morphological heterogeneity with distinct proliferative behaviors, transcriptional programs and lineage outcomes in neocortical development. Several aspects of our study deserve specific discussion: (1) the interplay between morphological heterogeneity and progenitor identity; (2) the relationship between bRG morphology and cell fate in neocortical development; and (3) the broader implications for brain development, evolution and pathologies.

### Morphological heterogeneity underlying progenitor diversity

bRG represent a major driver of cortical expansion in humans ^2,11,58^ yet the functional significance of their well-recognized morphological diversity ^7^ has remained largely unresolved. A central question is whether distinct bRG morphotypes identified in the human fetal neocortex ^18^ correspond to stable progenitor identities or transient cell states. While previous organoid studies have captured overall morphological dynamics of bRG ^20,59–61^, the morphotype-specific dynamics remained unresolved. Here we demonstrate that cortical CBOs preserve the full range of bRG morphotypes observed in fetal tissue (Figure 6A) and that individual morphotypes exhibit distinct morphodynamic profiles across the cell cycle. Monopolar and multipolar bRG inherit their morphological identity from mother cells, whereas bipolar, bifurcated and bifurcated bipolar morphotypes are more flexible and generate morphologically diverse progeny (Figure 6B). During interphase, most morphotypes remain stable except for multipolar cells, which show extensive remodeling. While BP morphological heterogeneity was reported across mammals ^17–21^, here we show that this diversity may be transcriptionally encoded and can emerge from specific progenitor subclasses during defined phases of the cell cycle (Figure 6A, B). These observations suggest that morphological stability and flexibility are intrinsic and morphotype-dependent properties, reflecting lineage diversity within the progenitor pool.

**Figure 6.**
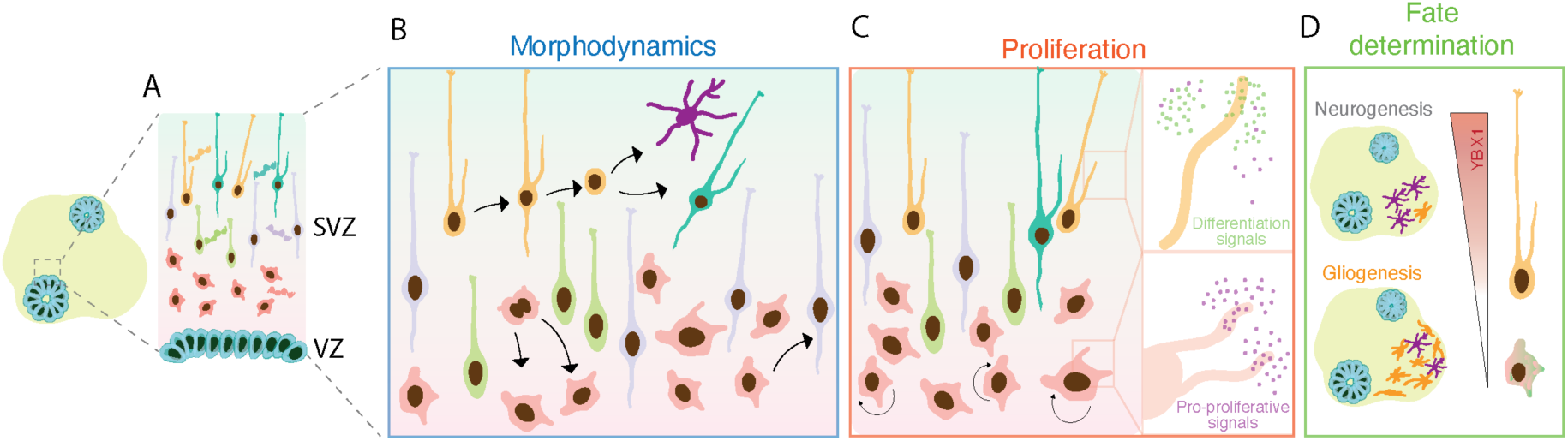
Human basal radial glia morphotypes exhibit different gene expression, proliferative capacities, morphodynamics and cell fate determination. **A)** Schematic representation of human CBOs containing morphologically diverse bRG with transcriptionally defined morphoclasses. **B)** Schematic model depicting how bRG morphotypes differ in their degree of structural remodeling across the cell cycle. **C)** Multipolar bRG are the most proliferative bRG morphotype. bRG proliferation is partially mediated by exposure to pro-proliferative vs differentiative signals. **D)** Diagram proposing a YBX1-associated neurogenic-to-gliogenic balance in a morphotype-specific manner.

Multipolar bRG also exhibit the greatest proliferative capacity and cell cycle re-entry rates, suggesting that structural flexibility may support continued progenitor activity (Figure 6C). Importantly, these SOX2+ HOPX+ multipolar cells were TBR2–, confirming they are *bona fide* bRG rather than bIPs (Figure S2). Their presence in fetal human neocortex ^18^ argues against an organoid artefact and allows us to hypothesize that multipolar bRG correspond to “transient bRG”, previously described in fetal macaque neocortex ^17^. Thus morphodynamics, particularly in multipolar bRG, may represent an evolutionary strategy to balance self-renewal with lineage diversification.

### bRG morphology and cell fate determination

Our data suggest that the morphological state of bRG correlates with their proliferative behavior and transcriptional identity. Spatial distribution of bRG morphotypes may also play a role here, as multipolar bRG cluster in the more apical portion of the SVZ, while bifurcated morphotypes reside in more basal positions. This allows us to hypothesize that the more apical location could be associated with the pro-proliferative niche, while basal positions could expose progenitors to local signals favoring neuronal differentiation. This agrees with our previous hypothesis linking progenitor morphology and proliferative capacity ^7^. In this model, growing additional cellular protrusions expose cells to extrinsic signals that can promote cell proliferation. We now extend this scenario by proposing that radial position within SVZ may be linked with the content of local niches and preferentially promote proliferation or differentiation.

CellShape-seq analysis shows that multipolar bRG display stem-like transcriptional profiles enriched for proliferation and gliogenesis pathways, whereas bifurcated bRG exhibit distinct signatures marked by the multifunctional gene expression regulator YBX1 ^47,48^. Through its roles in regulating mRNA stability, translation and modulating PRC2, YBX1 is known to regulate cortical development by promoting progenitor proliferation and neurogenesis ^53,54,62^. Here, we show that YBX1 has an essential role in maintaining bifurcated bRG identity and that upon its inhibition, bifurcated bRG are depleted, whereas multipolar bRG, that are the most proliferative, became a proportionally more abundant morphotype (Figure 6D). YBX1 inhibition further caused a marked decline of TBR2+ bIPs and SATB2+ late-born neurons, and a concurrent increase in astrocytic differentiation, indicating premature gliogenesis. Our findings suggest that YBX1 could potentially act upstream of the transition from neurogenesis to gliogenesis and when YBX1 is inhibited, the premature astrocytic differentiation occurs.

### Implications of bRG morphological diversity for brain development, evolution and disease

Our findings suggest that morphological heterogeneity within the bRG population is linked to lineage potential and may have contributed to neocortical expansion during evolution. Species with expanded and more complex neocortices, such as ferrets and macaques, exhibit greater bRG abundance and morphological diversity ^17–19^. These features are even more emphasized in human fetal brain development ^18,20^. This has led to a hypothesis that morphological complexity enables bRG to expand their surface and receive additional pro-proliferative signals from the local stem cell niche that in turn promote their proliferative capacity ^7^. The morphological flexibility observed in multipolar bRG could provide the functional plasticity required to generate the expanded neuronal output characteristic of gyrencephalic brains. This notion aligns with recent findings showing that molecular and metabolic pathways regulating bRG morphology, and hence their proliferative potential, have been evolutionarily adapted in modern humans compared to Neanderthals ^63^. Therefore, bRG morphology may be one of the features evolutionarily amplified in humans, supporting the development of a large, folded, and complex cortex.

Beyond evolution, the morphological and molecular diversity of bRG may represent points of vulnerability in neurodevelopmental disorders ^64–66^. Particularly interesting may be multipolar bRG as they upregulate various genes associated with intellectual disability and autism spectrum disorder (Figure S7F). Furthermore, dysregulation of YBX1 ^67,68^ or other gene expression regulators could disturb the balance between proliferation and differentiation, leading to aberrant gliogenesis or defective neuronal production. Linked to this, YBX1 was found to be significantly upregulated in high-grade gliomas ^69^ and involved in maintenance of glioblastoma stem cells ^70^, which in turn exhibit striking similarities to fetal bRG, both in terms of their molecular fingerprint and cell morphology ^71,72^. Understanding how morphology integrates various cell intrinsic programs with extrinsic cues will thus be critical for linking progenitor behavior to cortical malformations and neurological diseases.

## Acknowledgements

We are grateful to the National Facilities (NF) and services of Human Technopole (HT) for their outstanding support, notably the teams of the NF for Light Imaging, NF for Genomics and NF for Data Handling and Analysis. FKD thanks Chiara Ossola for technical assistance and MB thanks Cristina Cheroni for dataset preparation. We thank all the members of the Kalebic lab for helpful discussions. CB, MP and EC are Ph.D. students within the European School of Molecular Medicine (SEMM). Research in the Kalebic lab is supported by funds from HT and grants from AIRC (MFAG 2022 ID 27157) and Gilbert Family Foundation (#923004 and #925001) to NK.

## Author contributions

Conceptualization: FKD and NK; Methodology: FKD, CB, MB, MP, LW, EC, IB, DR, FC, AF, CP; Formal analysis: FKD, CB, MB, MP, RB; Investigation: FKD, CB, MP, LW; Data curation: MB; Writing – original draft: FKD and NK; Writing – review and editing: All authors; Visualization: FKD, MB, RB; Supervision: NK; Project administration: NK; Funding acquisition: NK.

## Resource availability

Further information and requests for resources and reagents should be directed to Nereo Kalebic).

Upon acceptance the raw data generated in this study will be available at the NCBI Sequence Read Archive (SRA).

## Key resource table

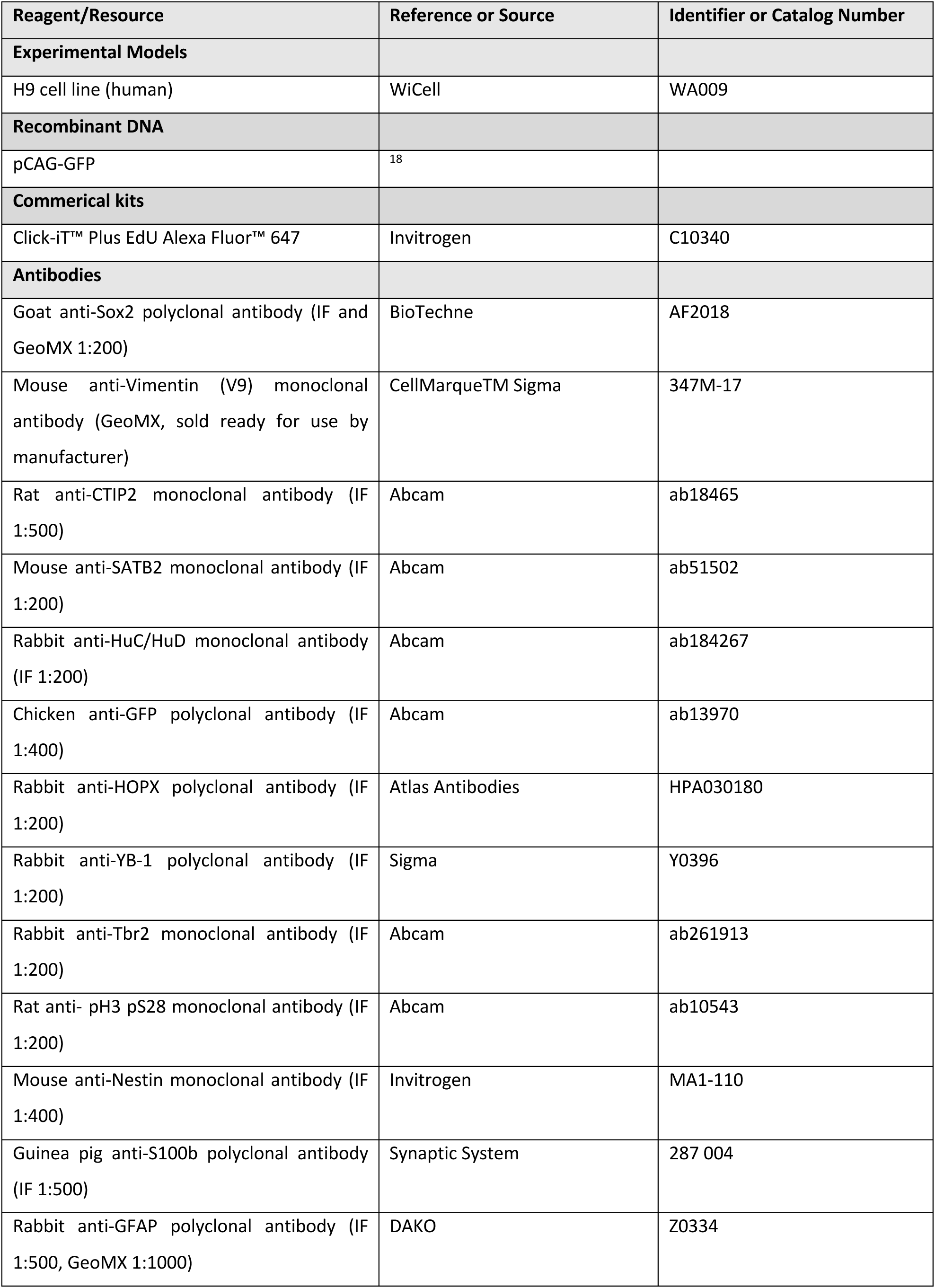
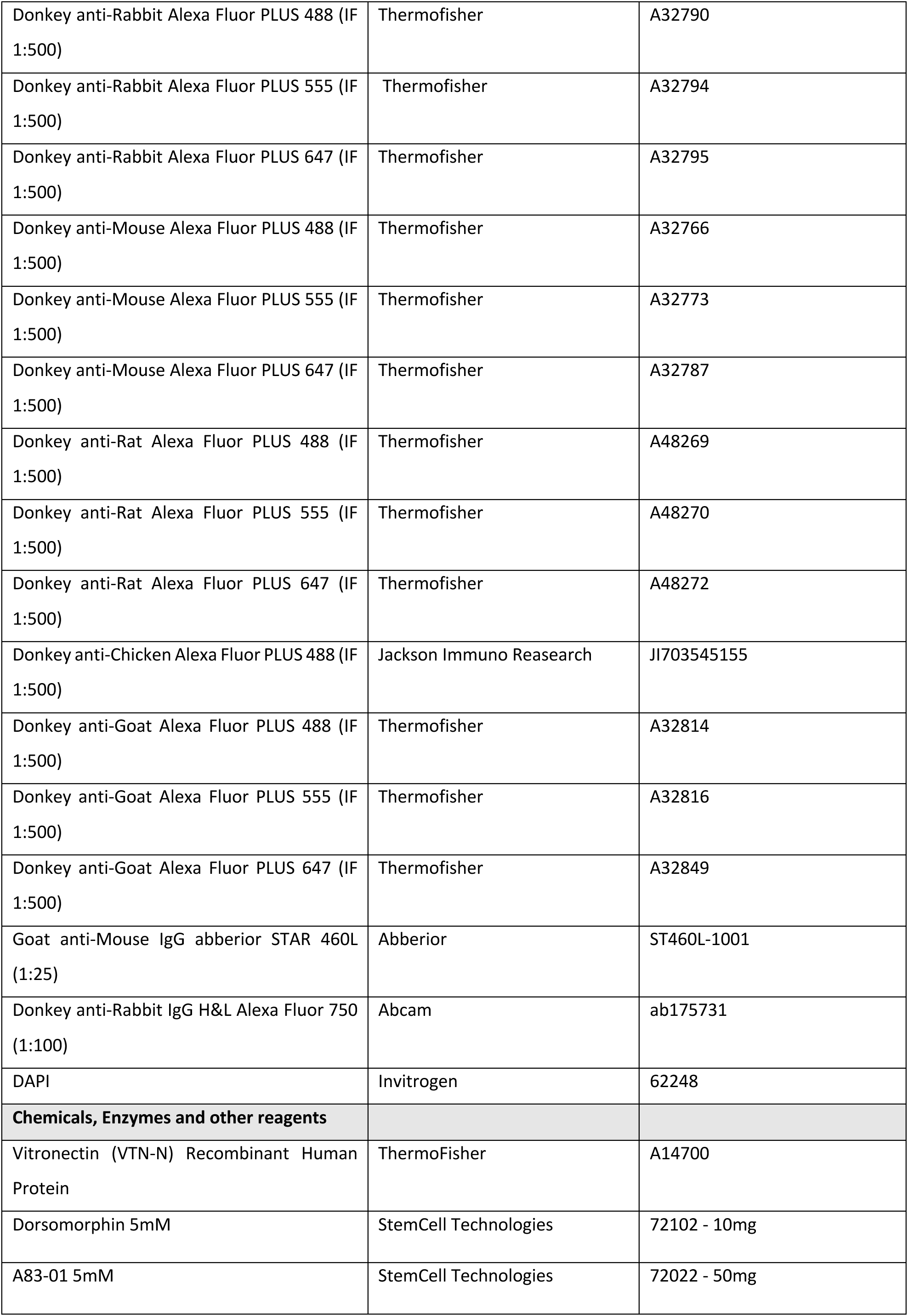
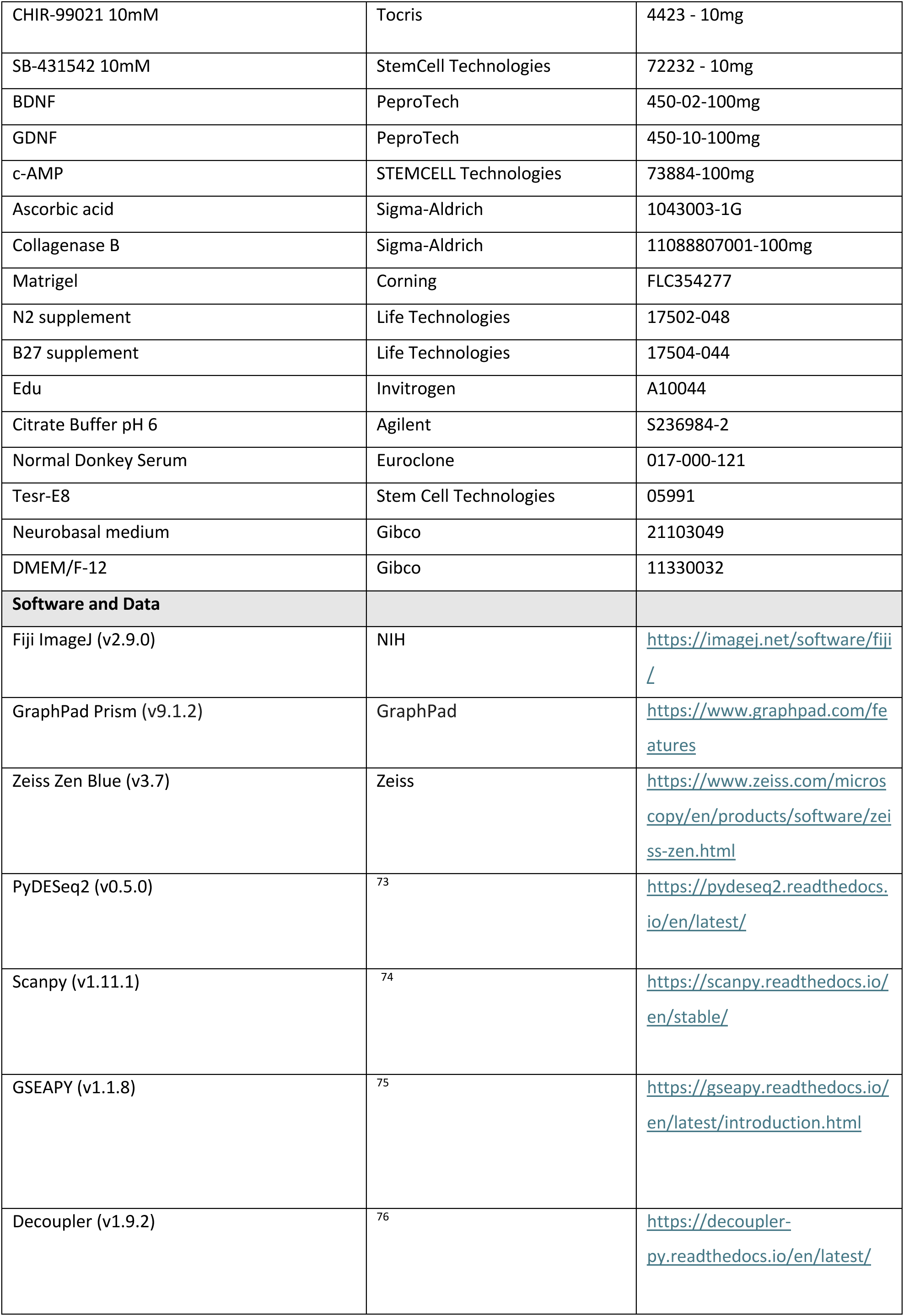
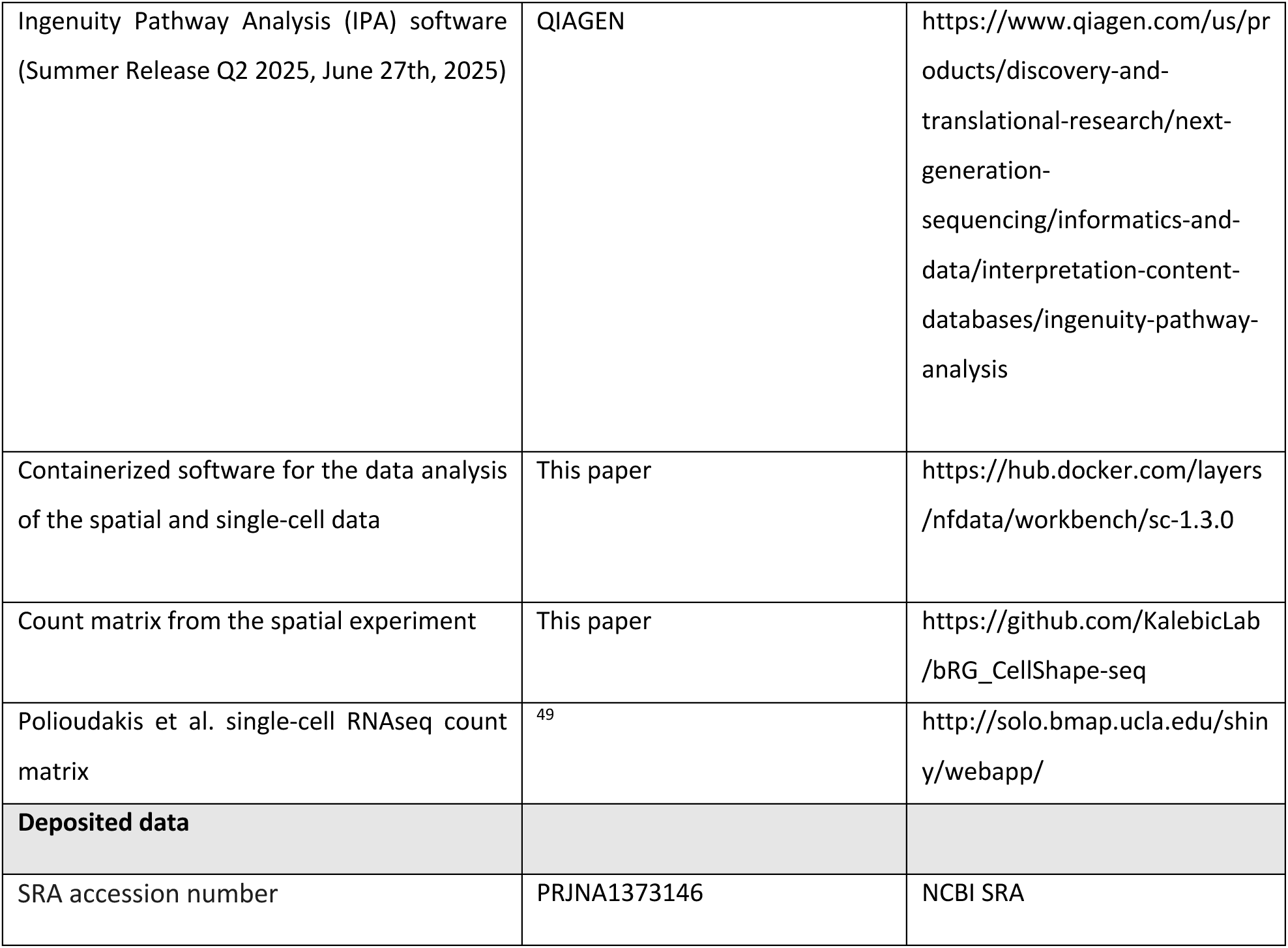

## Method details

### Stem cell and organoid culture

Human pluripotent stem cells (hPSCs) H9 were maintained on vitronectin-coated plates in a pluripotency-supporting medium (Tesr-E8) until colonies reached approximately 70–80% confluence, as previously described ^21^. Medium was changed daily.

Human cortical brain organoids (CBO) were generated following the protocol by ^34^. Briefly, for the organoid induction, hPSC colonies were detached from the plate by treatment with 1 mg/ml Collagenase Type IV for 10-15 min. Dissociated colonies were transferred to ultra-low attachment plates to form embryoid bodies (EBs). Neural induction was initiated by culturing the aggregates in medium containing 20% Knockout serum replacement (Gibco), 1X GlutaMAX (Gibco), 1X MEM-NEAA (Gibco), 1X Pen/Strep (Gibco), 2 µM Dorsomorphin (Stem cell technologies), 2 µM A83-01 (Stem cell technologies), 1X 2-Mercaptoethanol (Gibco) in DMEM/F12 (Gibco). On day 5 and day 6, half of the medium was replaced with induction medium consisting of DMEM/F12, 1X N2 Supplement, 1X Penicillin/Streptomycin, 1X Non-essential Amino Acids, 1X GlutaMax, 1 μM CHIR99021 and 1 μM SB-431542. On day 7, the EBs were embedded in droplets of Matrigel (Corning) to provide extracellular matrix support and cultured for an additional week to allow neuroepithelial formation. On day 14, the Matrigel-embedded CBOs were mechanically released and cultured with orbital shaking at 120 rpm in differentiation medium composed of DMEM/F12, 1X N2 and 1X B27 Supplements (Life Technologies), 1X Pen/Strep, 1X 2-Mercaptoethanol, 1X MEM-NEAA, 2.5 μg/ml Human Insulin (Sigma-Aldrich). From day 35 to day 70, the medium was supplemented with 1% Matrigel. On day 70, additional factors including ascorbic acid, cyclic AMP, BDNF, and GDNF were supplied to promote neuronal maturation. Medium changes were performed every 2 days. At indicated time points, CBOs were harvested for analyses. According to the protocol, the CBOs were sliced for maintenance, as follows. CBOs were embedded in 3% low melting point agarose (Invitrogen), dissolved in DMEM/F12 and sliced into 500 µm sections at 0.1 mm/s speed and 1 mm amplitude using a VT 1200S vibratome (Leica Microsystems) at days 45 and 72 were. Slices were cultured in differentiation medium supplemented with 5% FBS, EGF and FGF.

### CBO electroporation

CBO electroporation was performed as previously published ^21^. Briefly, CBOs were first embedded in 3% low-melting-point agarose and sectioned at a thickness of 200 µm using a vibratome (VT 1200S). pCAG_GFP (0.2 µg/µL in sterile PBS with 0.01% Fast Green (Sigma) for visualization) was microinjected into the ventricular-like cavities using a pulled glass micropipette under stereomicroscopic guidance. Electrical pulses were then applied with tweezer-type platinum electrodes (Manufacturer) connected to a square-wave electroporator (Manufacturer). Electrodes were positioned on opposite sides of the slice to direct current across the ventricular zone, and five pulses (50 V, 50 ms duration, 5 ms intervals) were delivered.

### Pharmacological treatment

CBOs were exposed to the small-molecule YBX1 inhibitor SU056 (100nM and 1 µM; ^55^, or DMSO (Vehicle). Two treatment paradigms were used: short-term inhibition (8 days) starting from week 14 COs and long term (chronic inhibition – 30 days) starting from week 12 CBOs. CBOs were maintained under standard differentiation conditions, and the inhibitor-containing medium was refreshed every 48 hours. At the end of the treatment, CBOs were fixed in 4% PFA and processed for microscopy as described below. Treatments were performed on at least 5 CBOs per condition per experiment from three independent differentiations.

### Sample preparation for microscopy and immunofluorescence

For the microscopy of fixed samples, CBOs fixed in 4% paraformaldehyde (PFA) for 30 min at room temperature were washed with PBS and cryoprotected in 30% sucrose overnight at 4 °C. Samples were embedded in OCT, sectioned at 20 µm on a cryostat, and mounted on Superfrost Plus slides. Next, samples were incubated in 10mM citrate buffer, pH 6 (70°C, 1 h) for antigen retrieval, washed in PBS and blocked/permeabilized in PBS containing 10% donkey serum and 0.3% Triton X-100 for 2 h at room temperature. Primary antibodies were applied overnight at 4°C, washed in PBS, and detected with Alexa Fluor–conjugated secondary antibodies (1:300, 2 h, room temperature). Nuclei were counterstained with DAPI (15 min, room temperature), and samples were mounted in Mowiol for imaging.

For live imaging, CBO slices after electroporation were transferred onto Millicell culture inserts (0.4 µm, PICMORG) and maintained at the air–liquid interface (ALI) in differentiation medium throughout the long-term live imaging, as previously published ^20,60^.

For immunofluorescence after live imaging, the samples were fixed overnight in 4% PFA at 4°C and processed for free-floating immunofluorescence. Briefly, the samples were transferred from the inserts into a 24 well plate and blocked/permeabilized as above. Primary antibodies were applied for 48h at 4°C and detected with Alexa Fluor–conjugated secondary antibodies as above. The cell fate was examined using the antibodies against HuC/HuD (neuronal marker) and SOX2 (progenitor marker).

### Microscopy

Confocal imaging of the fixed samples was performed using Zeiss LSM 980, Zeiss LSM 980 NLO or Leica Stellaris 8 point-scanning confocal microscopes. The Zeiss LSM 980 and LSM 980 NLO systems are mounted on Observer 7 inverted microscopes equipped with motorized stage and the following laser lines: 405 nm, 488 nm, 561 nm, 594 nm, 639 nm. The Leica Stellaris 8 system is mounted on a DMI8 inverted microscope equipped with a motorized stage and the 405 nm laser line plus the white light laser (WLL range: 440 nm - 790 nm). Images were acquired with a Plan-Apochromat 40x/1.4 NA oil immersion objective using Zen Blue 3.7 software (Zeiss) on the Zeiss confocal systems and with a Plan-Apochromat 40x/1.3 NA oil immersion objective using LAS X software (Leica) on the Leica system. For each organoid, five non-overlapping fields of view were selected across comparable radial positions to minimize sampling bias. For each field of view, z-stacks were acquired with a 0.2 µm z-step to analyze the whole sample volume. Free-floating sections immunostained for HuC/HuD and SOX2 were imaged on Zeiss LSM 980 confocal microscope in a drop of PBS in a glass-bottom dish.

For live imaging, GFP expression from electroporated slices was readily detected at 48 h after electroporation. Slices maintained in ALI culture were transferred on their Millicell inserts into glass-bottom imaging chambers and imaged using either Zeiss LSM 980 or Zeiss LSM 980 NLO confocal microscopes. During imaging, slices were incubated in a stage-top incubator at37°C in a 5% CO₂ atmosphere plus >90% humidity. Multiple imaging positions were defined across the SVZ-like regions of each slice. For each position, a z-stack spanning 80–100 µm was acquired with 5 µm z-step to image the entire SVZ and adjacent tissue while minimizing acquisition time. Time-lapses were performed with a time interval of 15 min for a total duration between 48 h and 60 h. The movies were acquired with a Plan-Apo 10x/0.45 dry objective using 488 nm laser line. The software used for all the acquisitions was Zen Blue 3.7. Image datasets were exported and analyzed using Fiji/ImageJ.

### CellShape-seq - sample preparation and sequencing

CellShape-seq was done as previously established with minor modifications ^33^. In brief, CBOs from three different batches were formalin fixed and paraffin embedded to be cut at 4 µm thickness at the microtome. Sample preparation was carried out following the GeoMx DSP Manual Slide Preparation (MAN-10150-06) and, for *in situ* hybridization, the Human Whole Transcriptome Atlas (GMX-RNA-NGS-HuWTA-4) was used. To perform manual segmentation of the bRG morphotypes, the morphology markers Vimentin (CellMarque, 347M-17), GFAP (Dako, Z0334), SOX2 (R&D, AF2018) were used along with the nuclear marker Syto13 (ThermoFisher, S7575). bRGs were defined as Vimentin+ SOX2+ GFAP+ cells in the CBO SVZ and they were grouped into three morphoclasses (monopolar+bipolar, multipolar and bifurcated).

Within each of the 18 CBOs, ROIs were drawn at the maximum size allowed (660x785 µm) and, within these regions, single cell segmentations for the three different morphoclasses were drawn creating three distinct AOI (areas of illumination) masks within each ROI. For the AOI segmentations, .tiff files of the ROI were extracted in Fiji ImageJ and manual cell segmentation was performed as described previously ^33^.

For library preparation and sequencing, oligos from AOIs were collected into a 96 well plate. They were then amplified and tagged with ROI-specific barcodes, adding sequencing adapters for subsequent high-throughput sequencing, following the protocol described in the GeoMX DSP NGS Readout User Manual (MAN-10153-06). Libraries were sequenced on a NextSeq2000 instrument (Illumina) generating per each AOI a number of read pairs derived by applying the following formula n. read pairs = total collection area (µm2) * sequencing depth factor (equal to 100 for the WTA panel).

### Image analysis – cell morphology and cell counts

Time-lapse confocal datasets were analyzed to determine bRG morphology and its inheritance. To focus the analysis on progenitor cells, only GFP+ cells that underwent at least one division during the imaging period were analysed. Morphology was annotated frame by frame and cells were classified as monopolar, bipolar, bifurcated, bifurcated bipolar, or multipolar according to the number and orientation of processes. For inheritance analysis, the morphology of each mother cell was determined at the last frame before mitosis and compared directly to that of its daughter cells in the first frame after cytokinesis. For interphase morphology, cell shape was assessed either between the start of imaging and the last frame before the mitosis, or between the first frame after the cytokinesis and either the end of imaging or the start of the next division. All morphotype annotations were performed manually on 3D stacks using Fiji/ImageJ and analyses were restricted to divisions in which complete lineages could be tracked throughout the imaging period.

For fixed samples individual optical sections were used to identify bRG based on SOX2 and HOPX expression. Cell morphotype (monopolar, bipolar, multipolar, bifurcated and bifurcated-bipolar) was annotated according to the number and orientation of primary processes. Only cells with clearly traceable processes throughout the Z-stack were included in the analysis. Morphotype quantification was performed manually in Fiji, and counts were normalized per field of view or per CBO as indicated.

Quantification of cell counts for all experiments was performed manually in Fiji, and counts were normalized per field of view or per CBO as indicated. For cell fate assigning, imaging positions recorded during live acquisition were manually re-identified by aligning the final live frame with post-fixation confocal image using GFP signal and structural landmarks. Daughter cells observed in the last live frame were then matched to their post hoc immunostaining profile, allowing assignment of cell fate as neurons (HuC/HuD+) or progenitors (SOX2+).

For the subcellular localization of YBX1, immunofluorescence images were analysed manually to determine the presence of YBX1 within the cell body, nucleus, and cellular processes. For each compartment, morphotypes were scored as positive or negative based on detectable signal above background. Quantification shows the percentage of morphotypes positive for each compartment in control and treated organoids.

All image analysis data were analyzed in GraphPad Prism. We used two-way ANOVA with post-hoc corrections where appropriate or Mann Whitney U-test. Error bars at all graphs represent mean ± s.e.m.; significance thresholds: *p*<0.05, p<0.01, *p*<0.001, *p*<0.0001. The number of independent organoid differentiations, CBOs or individual cells is reported in each figure legend.

### CellShape-seq dataset analysis

The analysis of the CellShape-seq dataset was performed as previously established ^33^. Briefly, raw sequencing reads were processed using the GeoMx NGS Pipeline (NanoString, version 2.3.3.10) for alignment, deduplication, and quantification of the probes from the GeoMx Human Whole Transcriptome Atlas. Differential expression analysis was conducted using PyDESeq2 v0.5.0 ^73^ on raw counts. A model was fitted, incorporating the area of interest (AOI) morphoclass and the experimental GeoMX run as covariates. Gene expression contrasts were computed for each morphoclass (multipolar, bifurcated, and monopolar+bipolar) against each other. Genes with an adjusted p-value < 0.05 were considered significantly differentially expressed. Such genes were functionally characterized through over-representation analysis of Biological Process Gene Ontology (GO) terms (2024.1.Hs version from MSigDB ^77^. Enrichment analysis was performed using the enrichr function from gseapy ^75^.

### Functional enrichment analysis of CellShape-seq dataset

Functional enrichment analysis of differentially expressed genes (DEGs) was performed using QIAGEN Ingenuity Pathway Analysis (IPA) ^78^. A dataset containing gene identifiers and corresponding expression values from multipolar and monopolar+bipolar versus bifurcated morphoclasses was uploaded to the IPA platform. Each gene identifier was mapped to its corresponding entity within the QIAGEN Knowledge Base. Significantly perturbed molecules (FDR < 0.05) were then superimposed onto a global molecular network curated from the QIAGEN Knowledge Base. Biological function enrichment was assessed using a right-tailed Fisher’s Exact Test to calculate p-values, estimating the likelihood that the observed associations between DEGs and specific biological functions or diseases occurred by chance. Upstream Regulator Analysis was conducted to identify potential transcriptional regulators responsible for the observed expression changes and to predict their activation state. A Z-score was computed to infer the directionality of regulation (activation or inhibition).

### Mapping CellShape-seq dataset to a fetal scRNA-seq dataset

The dataset from ^49^ was obtained from the CoDEx Viewer platform [website: http://solo.bmap.ucla.edu/shiny/webapp/]. To discard low-quality cells and doublets, barcodes were filtered according to the following criteria: (1) number of detected genes lower than 500 or higher than 5,000, (2) number of counts lower than 750 or higher than 10,000; (3) % of mitochondrial counts higher than 5%, (4) % ribosomal protein counts higher than 20%. Genes expressed in less than 0.25% of the cells were also filtered out. To focus on the cell populations of interest, the dataset was then subset to the cells annotated as oRG (Outer Radial Glia), OPC (Oligodendrocyte Progenitor Cells), IP (Intermediate Progenitors), PgS (Progenitors in S phase), and PgG2M (Progenitors in G2/M phase) by the authors. The count matrix was normalized by scaling the total cell counts to 10,000, excluding the counts of highly expressed genes. Then a reduced representation of the data (principal component analysis, UMAP) was built on the highly variable genes, following the standard Scanpy ^74^ workflow. We then classified *bona fide* multipolar and monopolar+bipolar cells in the dataset, by projecting the transcriptomic signatures obtained from the GeoMX differential expression analysis. From the genes differentially expressed in the multipolar vs bifurcated and the monopolar+bipolar vs bifurcated differential analysis, we derived two gene signatures selecting the genes that were identified as up-regulated (log₂ fold-change > 0). The resulting sets of multipolar-upregulated and monopolar+bipolar-upregulated genes were used as signatures for downstream enrichment analysis.

To quantify the activity of these signatures in single cells, we implemented a rank-based single-cell enrichment test. For each cell in the ^49^ dataset, all genes were ranked according to their standardized expression (z-scored values). For each of the two signatures, we compared the distribution of ranks of the signature genes to that of all remaining genes using a one-sided Mann–Whitney U test. For each cell, two outputs were then computed: (i) a normalized enrichment score (NES), defined as the Mann–Whitney U statistic scaled by nm, with n being the number of genes in the gene list and m being the number of background genes, and (ii) an associated p-value. This procedure produces a per-cell enrichment measure analogous to a single-sample GSEA score based on rank separation. The resulting enrichment scores were then used to classify *bona fide* multipolar and monopolar+bipolar cells within the ^49^ dataset. A fixed enrichment threshold was applied: cells with NES > 0.5 for the multipolar signature were labeled as bona-fide multipolar cells, and analogously, cells with NES > 0.5 for the radial signature were labeled as bona-fide radial cells.

To identify a subset of progenitor cells enriched in activity of YBX1, we first inferred the regulatory activity at single-cell resolution for the progenitor cells from the dataset of ^49^. Regulatory activity of transcription factors was inferred using the ulm function implemented in the decoupler framework ^76^. This Univariate Linear Model (ULM) method computes transcription factor activity scores by modeling gene expression data with prior regulatory knowledge. As prior information, we used the regulons defined in the CollecTRI network ^79^, which provides curated transcription factor–target interactions. ULM quantifies regulator activity by fitting a simple linear model in which the expression of each target gene is explained by the regulator’s predicted influence. This produces a continuous activity score for each cell that reflects the predicted regulatory influence of each transcription factor. To distinguish between YBX1-activated and non-activated cells, we modeled the distribution of inferred YBX1 activity values using a two-component Gaussian mixture model (GMM). The model was fitted to the continuous activity scores to decompose the distribution into two Gaussian components representing low-activity and high-activity cell populations. The intersection point between the probability density functions of the two fitted components was computed and selected as the optimal threshold for separating the two populations. The resulting intersection value (activity = 1.40) was used as the cutoff to define a subset enriched in YBX1-activated cells. All cells with inferred activity scores above this threshold were classified as YBX1+ enriched for downstream analyses.

