## Supplemental information for "Human basal radial glia morphotypes are transcriptionally distinct and exhibit different cell fate determination"

Kaluthantrige Don et al.

### Supplemental figures and legends

### Figure S1

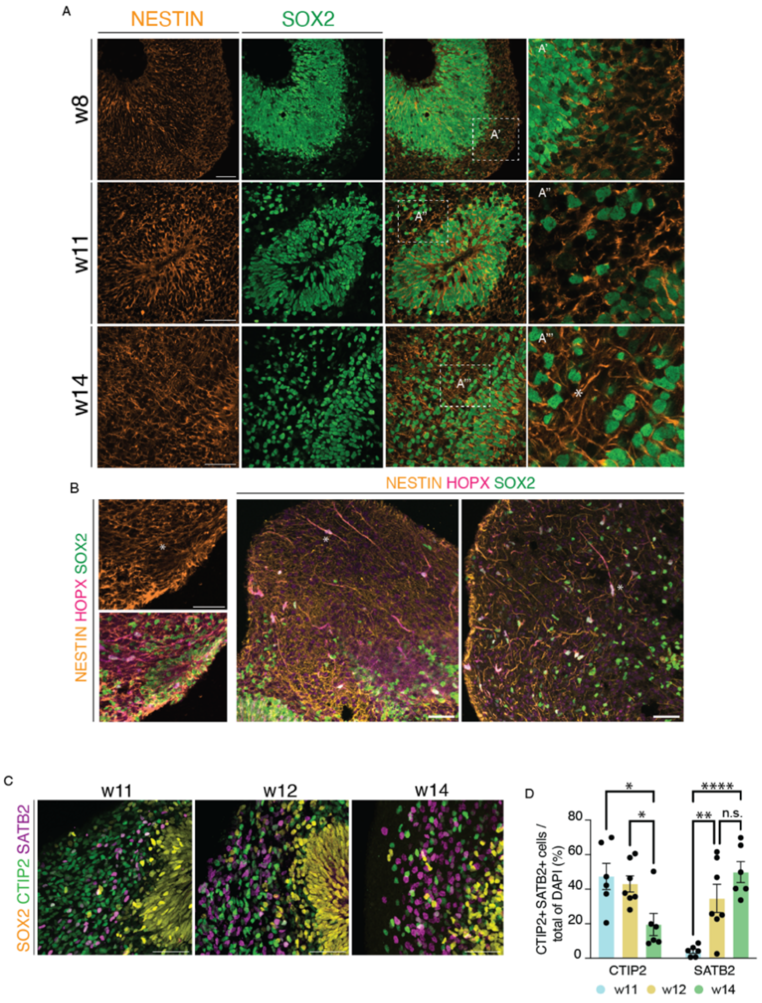


**Figure S1. Basal fibers originated in the SVZ and the production of SATB2+ late-born neurons both start after the onset of bRG.**

**A)** IF for nestin and SOX2 of CBOs w8, w11 and w14**.** Dashed boxes, areas magnified on the right. Scale bars, 50 µm. **B)** IF for SOX2 of w12 CBOs, nestin and HOPX, showing co-localization of nestin+ and HOPX+ fibers in the SVZ. Scale bars, 50 µm. Asterisk, start of the nestin+ fiber in the SVZ. **C)** IF for SOX2, CTIP2 and SATB2 in w11, w12 and w14 CBOs. Scale bars, 50 µm. **D)** Quantification of CTIP2+ and SATB2+ cells in w11, w12 and w14 CBOs. Error bars, SEM; N= 4 ; Mann-Whitney u-test, * $p< 0.05$; ** $p< 0.01$; **** $p< 0.0001$.

### Figure S2

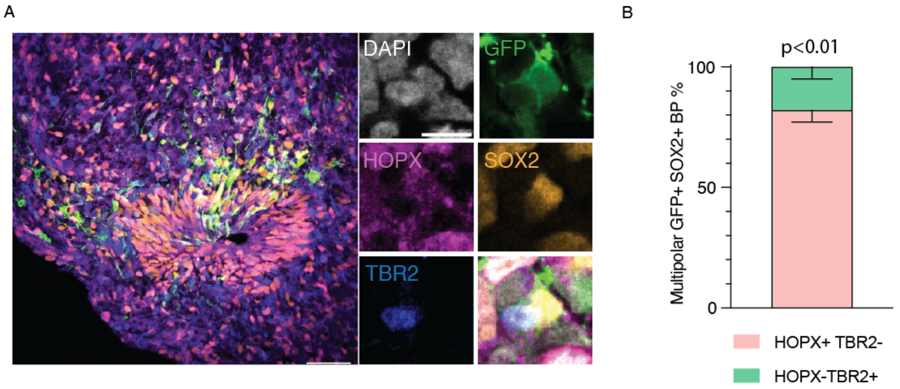


**Figure S2. Multipolar bRG are TBR2–.**

**A)** IF for GFP, SOX2, HOPX, and TBR2, along with DAPI staining, upon the electroporation of CBOs with GFP to visualized BP shape. Left, overview. Right, High magnification of a representative multipolar bRG. Scale bars, 50 µm (overview); 10 µm (high magnification). **B)** Quantification of multipolar BPs. Error bars, SEM; N = 41 cells from 3 different CBO differentiations; Mann-Whitney u-test, ** p< 0.01. Note that all the multipolar SOX2+ HOPX+ bRG, identified in Figure 1, do not express TBR2, a marker of basal intermediate progenitors (bIPs), which are also multipolar. Therefore, multipolar SOX2+ HOPX+ can be considered as *bona-fide* bRG.

### Figure S3

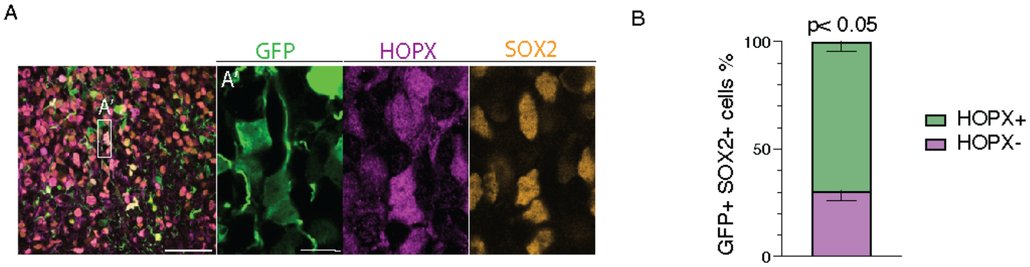


**Figure S3. Electroporated GFP+ cells in the SVZ are predominantly HOPX+ and SOX2+.**

**A)** IF for GFP, SOX2 and HOPX, upon the electroporation of CBOs prior to live imaging. Box, area magnified in the 3 insets on the right. Scale bars, 50 µm (overview); 10 µm (high magnification). **B)** Quantification of GFP+ SOX2+ electroporated cells in the SVZ based on HOPX+ positivity. Error bars, SEM; N= 4; Mann-Whitney u-test, * p< 0.05. Note that for the live imaging we analyzed only the cells that underwent cell division, while this quantification shows all the GFP+ cells. This implies that more than 70% of imaged cells are bRG.

### Figure S4

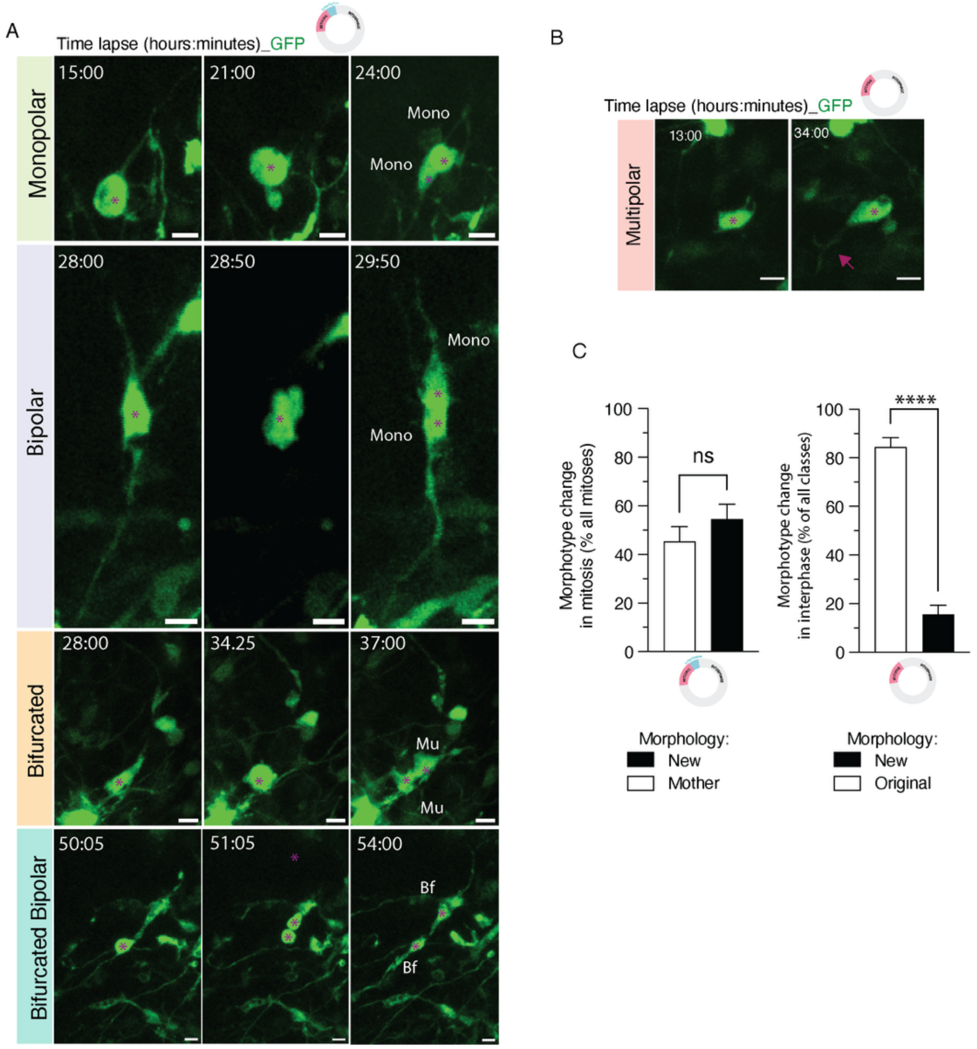


**Figure S4. Morphodynamics of bRG.**

**A)** Additional examples (to Figure 2C, E) of time-lapse sequences of individual bRG morphotypes undergoing division along with the morphology of their daughter cells. Asterisks, cell body; Mono, monopolar; Mu, multipolar; Bf, bifurcated. Scale bars, 10 µm. **B)** Additional example (to Figure 2F) of time-lapse sequence of a multipolar bRG elongating in interphase. Scale bars, 10 µm. **C)** Quantification of morphotype change in mitosis (left) and interphase (right) showing a high degree of morphological changes in mitosis and predominantly morphological stability in interphase. Error bars, SEM; N = 134 (mitosis) and 130 (interphase) bRG from 11 independent experiments; Mann-Whitney u-test, **** $p < 0.0001$; ns, statistically not significant. See also Movies S7-S11.

### Figure S5

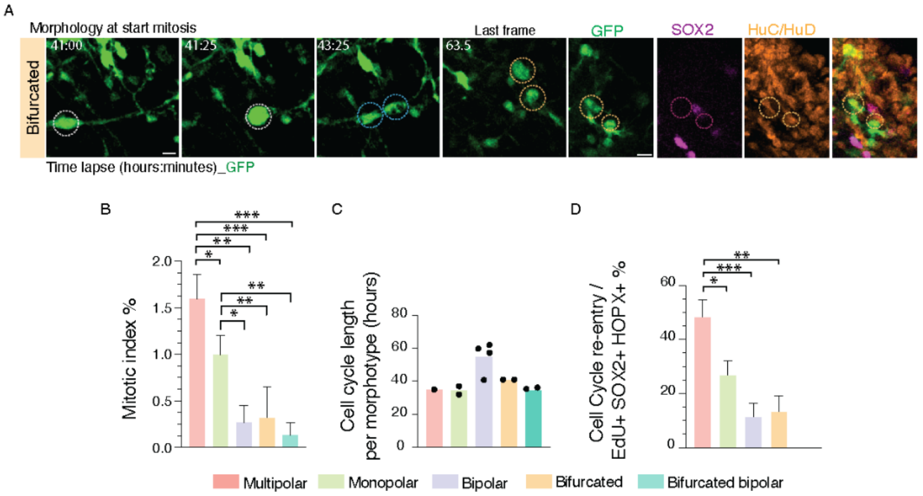


**Figure S5. bRG morphotypes exhibit different proliferative capacities.**

**A**) Representative images of bifurcated bRG dividing into two HuC/HuD+ neurons. Live imaging and fate mapping were performed as described in Figure 3. Scale bars, 10 µm. **B, D)** Quantification of the mitotic index (PH3+ / DAPI) (B) and proliferative capacity (Ki67+ EdU+ / EdU+) (D) across SOX2+ HOPX+ bRG morphotypes (irrespective of the relative morphotype abundance). Error bars, SEM; N = 58 (B) and 308 (D); Mann-Whitney test, * p< 0.05; ** p< 0.01; *** p< 0.001; **C)** Quantification of cell cycle length per morphotype for cells that underwent two cell cycles within the imaging timeframe. Each dot represents a cell.

### Figure S6

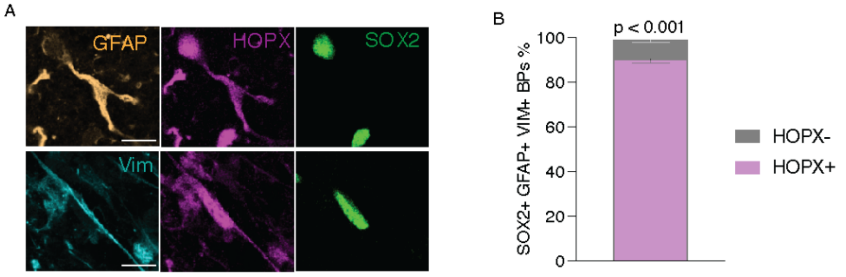


**Figure S6. GFAP+ Vim+ SOX2+ bRG profiled by CellShape-seq are predominantly HOPX+.**

**A**) Representative images of bRG upon IF for HOPX, SOX2, Vimentin and GFAP. Scale bars, 10 µm. **B)** Quantification of SOX2+ GFAP+ Vimentin+ BPs for HOPX positivity. Error bars, SEM; N = 306 cells from 9 independent differentiations; Mann-Whitney test, **** p< 0.0001.

**Figure S7**

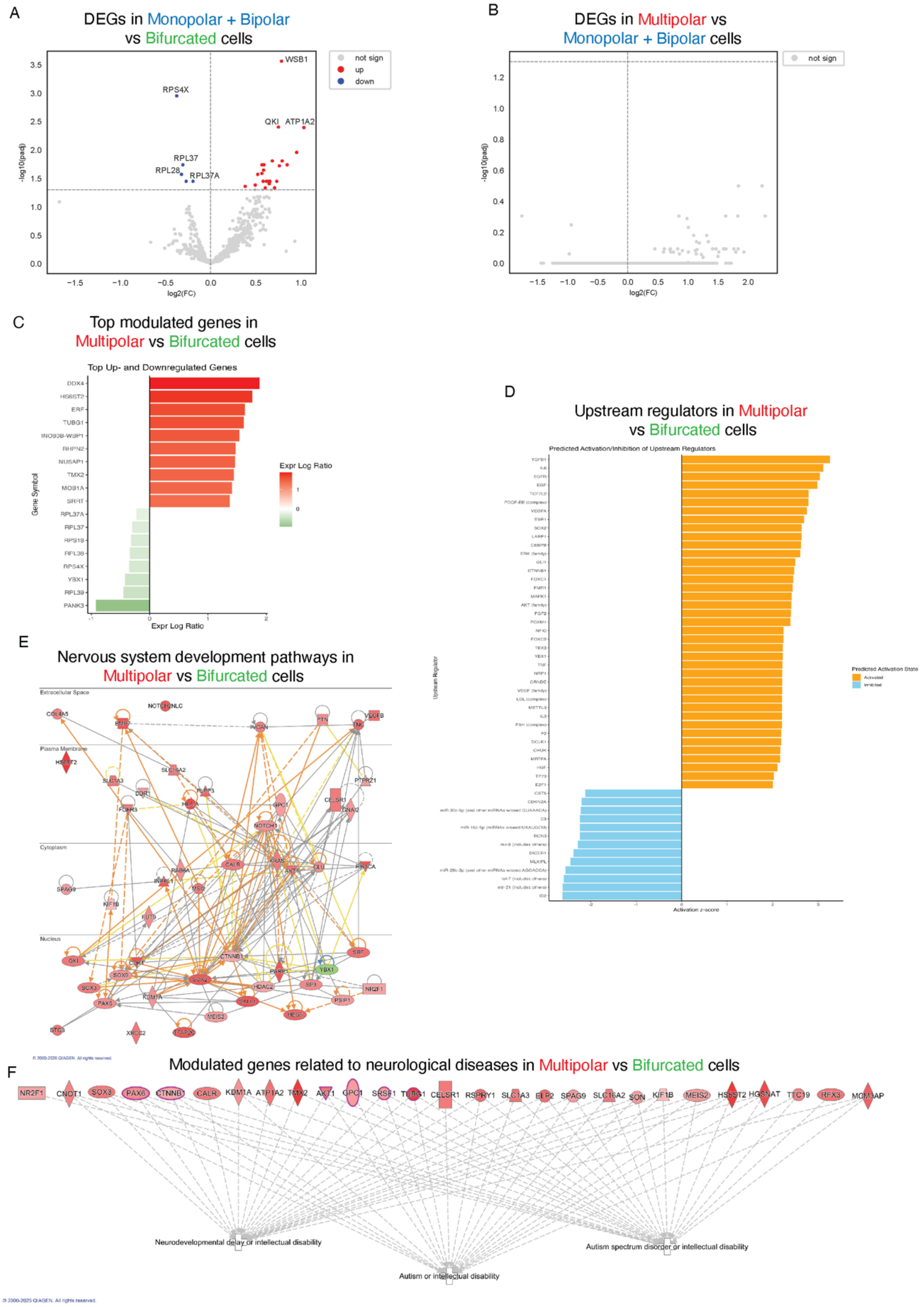

© 2000-2025 QIAGEN. All rights reserved.

**Figure S7. Modulators of multipolar and bifurcated bRG.**

**A-B**) Volcano plot showing the results of a differential expression analysis of monopolar+bipolar vs bifurcated bRG (A) and multipolar vs monopolar+bipolar bRG (B), The x-axis, LFC; y-axis, negative log10 of the adjusted p-value. Red dots, significantly up-regulated genes (LFC > 0 and adjusted p-value < 0.05); blue dots, significantly down-regulated genes (LFC < 0 and adjusted p-value < 0.05). **C)** Bar plots showing the Top Modulated Genes for the comparison of Multipolar vs Bifurcated cells. The graph was generated using QIAGEN Ingenuity Pathway Analysis (IPA). The intensity of the bar plot coloring corresponds to log2FC values, with up-regulated genes shown in red and down-regulated genes indicated in green. **D)** Bar plots illustrating the top Upstream Regulators identified by QIAGEN IPA's Upstream Regulator Analysis for the comparison between Multipolar and Bifurcated bRG. An upstream regulator is any molecule influencing expression, transcription, or phosphorylation of others. The analysis predicts which regulators are activated or inhibited based on observed gene expression changes. IPA compares the direction of change in known targets to literature expectations and uses a z-score algorithm for predictions. In the plots, orange bars indicate predicted activation, and light blue bars indicate predicted inhibition, with color intensity reflecting z-score magnitude. **E)** Graphical representation of the molecular relationships between modulated genes related to nervous system development in multipolar cells vs bifurcated bRG. Genes are depicted as nodes with edges illustrating biological relationships, solid lines for direct interactions and dashed lines for indirect ones. All edges are supported by at least one reference from the literature, textbooks, or canonical information in the QIAGEN Knowledge Base. Nodes are displayed in various shapes representing the functional class of the gene product. The intensity of the node color indicates the degree of up-(red) or down-(green) regulation. Edge colors convey predicted effects: orange suggests activation, blue indicates inhibition, yellow indicates possible inconsistencies with the downstream molecule's state, and grey denotes an undetermined effect based on current literature. **F)** Modulation of genes associated with neurological disorders in multipolar vs bifurcated bRG. Genes significantly modulated (pV < 0.05) and associated with autism spectrum disorders, intellectual disability, and other neurodevelopmental conditions are shown as nodes with distinct shapes indicating the functional class of the encoded protein. The intensity of the red color reflects the degree of upregulation. White crosses indicate the associated biological functions. The graph was generated using QIAGEN IPA. All edges are supported by at least one reference from the literature, textbooks, or canonical information in the QIAGEN Knowledge Base. Among the most upregulated genes are *TMX2*, whose dysfunction leads to severe brain developmental abnormalities (Vandervore et al. 2019); *HGNAT*, with variants linked to abnormal facial morphology and intellectual disability (Mudassir et al. 2024); *HS6ST2*, associated with X-linked intellectual disability (XLID) (Paganini et al. 2019); and *TUBG1*, whose alterations cause tubulinopathies characterized by brain malformations, microcephaly, early-onset epilepsy, and motor impairment (Poirier et al. 2013); NR2F1, associated with intellectual disability (Bosch et al. 2014); PAX6, whose deletion is associated with WAGR syndrome, characterized by intellectual disability (Wang et al. 2022); SOX3, critical for the development of the pituitary gland, brain, and face; mutations can lead to hypopituitarism, intellectual disability, and craniofacial abnormalities (Li et al. 2022).

**Figure S8**

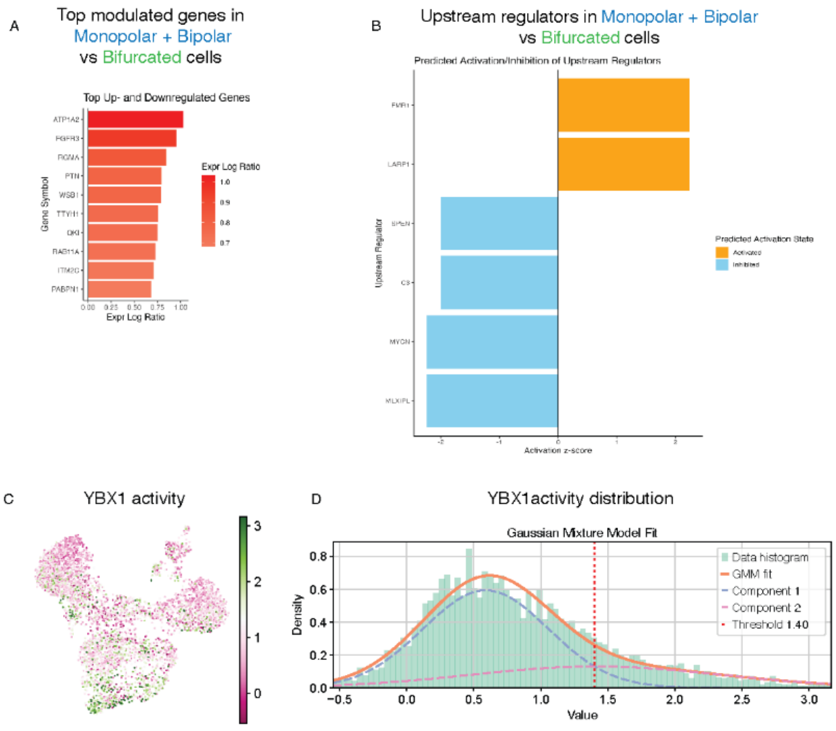


**Figure S8. Modulators of monopolar+bipolar bRG and the inferred activity of bifurcated bRG-enriched YBX1.**

**A)** Bar plots showing the Top Modulated Genes for the comparison of monopolar+bipolar vs bifurcated bRG. The graph was generated using QIAGEN IPA. The intensity of the bar plot coloring corresponds to log2FC values, with up-regulated genes shown in red. **B)** Bar plots illustrating the top Upstream Regulators identified by QIAGEN IPA's Upstream Regulator Analysis for the comparison between monopolar+bipolar vs bifurcated cells. See Figure S7D for details of annotation. **C)** Inferred activity of the YBX1. UMAP plot of progenitor cells from the dataset of Polioudakis et al. 2019, showing the inferred YBX1 activity values computed as described in Methods. **D)** Identification of a threshold (dotted red line) to define a YBX1+-enriched subset. The distribution of inferred YBX1 activity values from panel C is modeled using a two-component Gaussian mixture model, separating the component corresponding to YBX1-activated cells (blue dashed line) from that of non-activated cells (pink dashed line). The intersection point of the two modeled distributions (activity value = 1.40) is identified as the optimal threshold for defining a subset enriched in YBX1-activated cells. The cells exceeding this value are classified as YBX1+ enriched.

**Figure S9**

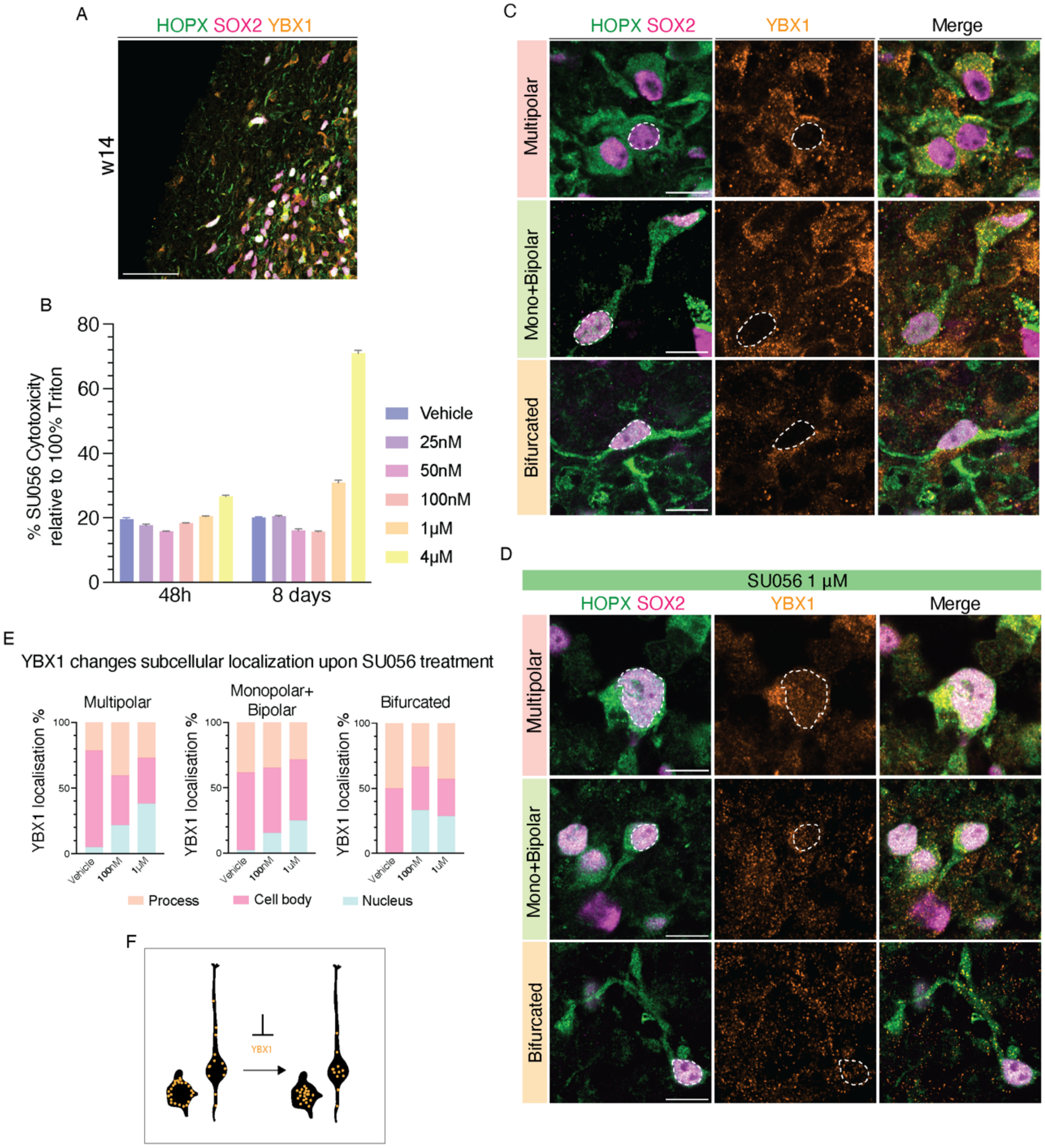


**Figure S9. YBX1 inhibitor SU056 changes subcellular localization of YBX1 in bRG.**

**A)** IF for HOPX, SOX2 and YBX1 of w14 CBOs. Overview. Scale bar, 50 μM. **B)** Dose response bar plot of YBX1 inhibitor SU056 after 48h and 8 days of treatment of CBOs, calculated relative to the negative control. **C, D)** IF for HOPX, SOX2 and YBX1 of bRG in w14 CBOs without (C) or upon 8 days of SU056 (YBX1 inhibitor) treatment (D). Scale bars, 10 μm. Dashed area, cell nuclei. **E)** Increase of YBX1 nuclear localization (blue bar) upon YBX1 inhibition. N = 131 (vehicle), 98 (100nM) and 105 (1μM) bRG. **F)** Schematic representation showing a bipolar and a multipolar bRG expressing YBX1 (yellow dots) in the processes and cell body. Upon YBX1 inhibition, YBX1 expression shifts in the nucleus.

**Figure S10**

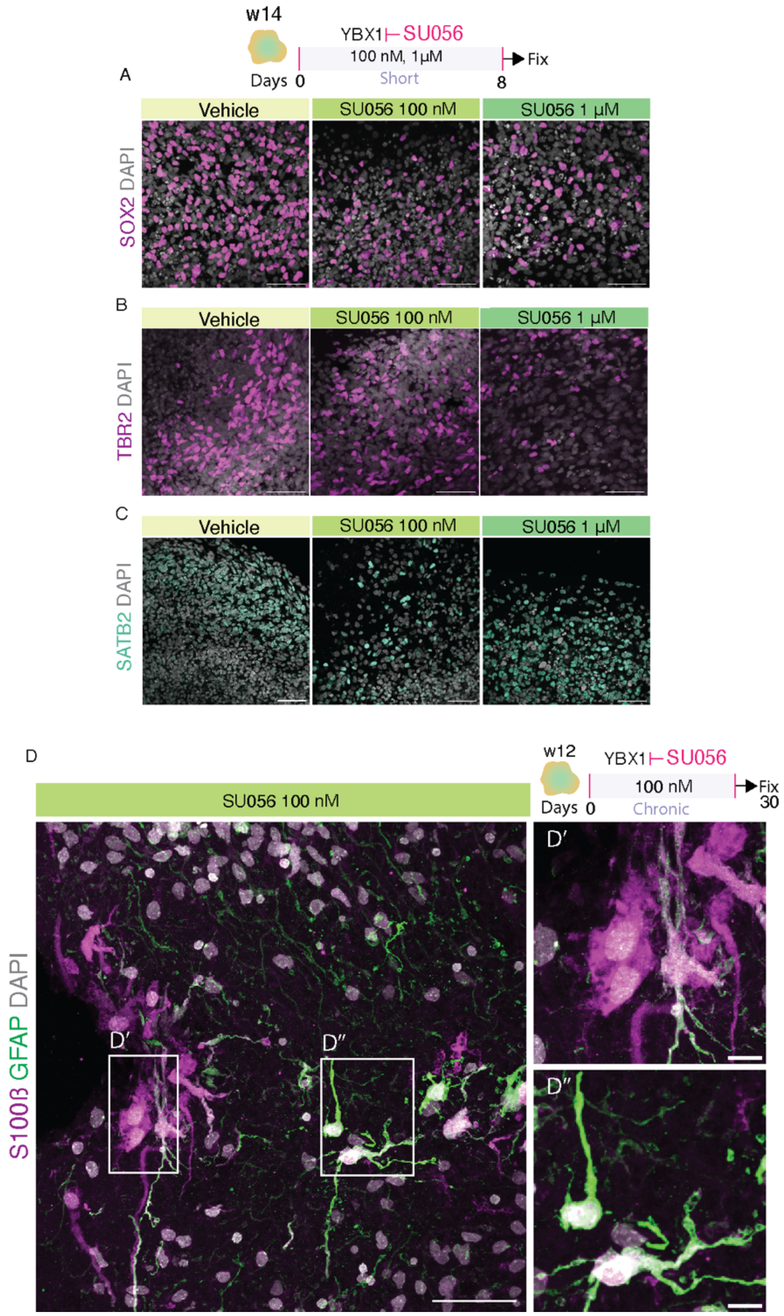


**Figure S10. YBX1 inhibition reduces abundance of progenitors and neurons and increase astrocytes.**

**A-C)** Example IF for SOX2 (A), TBR2 (B) and SATB2 (C) along with DAPI staining upon SU056 treatment. Scale bar, 50 µm. **D)** Long term inhibition of YBX1. Upper right, schematic representation of CBOs treated for 30 days (chronic) with SU056 (w12-16). Lower, IF for astrocytic markers S100ß and GFAP in CBOs upon chronic YBX1 inhibition. Boxes 1 and 2, areas magnified in insets on the right. Insets, higher magnification of astrocytes. Scale bars, 50 µm (overview); 10 µm (inset).

Li, Jing, Yuxia Zhong, Tao Guo, Yerong Yu, and Jianwei Li. 2022. "Case Report: A Novel Point Mutation of SOX3 in a Subject with Growth Hormone Deficiency, Hypogonadotrophic Hypogonadism, and Borderline Intellectual Disability." *Frontiers in Endocrinology* 13 (February): 810375.

Mudassir, Behjat Ul, Mujaddid Mudassir, Jamal B. Williams, and Zehra Agha. 2024. "Genetic Heterogeneity in Four Probands Reveals HGSNAT, KDM6B, LMNA and WFS1 Related Neurodevelopmental Disorders." *Biomedicines* 12 (12): 2736.

Paganini, Leda, Loubna A. Hadi, Massimiliano Chetta, Davide Rovina, Laura Fontana, Patrizia Colapietro, Eleonora Bonaparte, et al. 2019. "A HS6ST2 Gene Variant Associated with X-Linked Intellectual Disability and Severe Myopia in Two Male Twins." *Clinical Genetics* 95 (3): 368–74.

Poirier, Karine, Nicolas Lebrun, Loic Broix, Guoling Tian, Yoann Saillour, Cécile Boscheron, Elena Parrini, et al. 2013. "Mutations in TUBG1, DYNC1H1, KIF5C and KIF2A Cause Malformations of Cortical Development and Microcephaly." *Nature Genetics* 45 (6): 639–47.

Vandervore, Laura V., Rachel Schot, Chiara Milanese, Daphne J. Smits, Esmee Kasteleijn, Andrew E. Fry, Daniela T. Pilz, et al. 2019. "TMX2 Is a Crucial Regulator of Cellular Redox State, and Its Dysfunction Causes Severe Brain Developmental Abnormalities." *The American Journal of Human Genetics* 105 (6): 1126–47.

Wang, Qiwei, Xulin Zhang, Tingfeng Qin, Dongni Wang, Xiaoshan Lin, Yuanyuan Zhu, Haowen Tan, et al. 2022. "Unusual Presentation in WAGR Syndrome: Expanding the Phenotypic and Genotypic Spectrum of the Diseases." *Genes* 13 (8): 1431.
